# Mitochondria- and ER-associated actin are required for mitochondrial fusion

**DOI:** 10.1101/2023.06.13.544768

**Authors:** Priya Gatti, Cara Schiavon, Julien Cicero, Uri Manor, Marc Germain

## Abstract

Mitochondria play a crucial role in the regulation of cellular metabolism and signalling. Mitochondrial activity is modulated by the processes of mitochondrial fission and fusion, which are required to properly balance respiratory and metabolic functions, transfer material between mitochondria, and remove defective mitochondria. Mitochondrial fission occurs at sites of contact between the endoplasmic reticulum (ER) and mitochondria, and is dependent on the formation of actin filaments that drive mitochondrial constriction and the recruitment and activation of the dynamin-related GTPase fission protein DRP1. The requirement for mitochondria- and ER-associated actin filaments in mitochondrial fission remains unclear, and the role of actin in mitochondrial fusion remains entirely unexplored. Here we show that preventing the formation of actin filaments on either mitochondria or the ER disrupts both mitochondrial fission and fusion. We show that fusion but not fission is dependent on Arp2/3, whereas both fission and fusion are dependent on INF2 formin-dependent actin polymerization. We also show that mitochondria-associated actin marks fusion sites prior to the dynamin family GTPase fusion protein MFN2. Together, our work introduces a novel method for perturbing organelle-associated actin filaments, and demonstrates a previously unknown role for actin in mitochondrial fusion.

## Introduction

Mitochondria play a crucial role in the regulation of cellular metabolism and signalling, thereby controlling key cellular processes including apoptosis, cell fate decisions, and inflammation. These activities are regulated by mitochondrial dynamics, the process of fusion and fission which controls mitochondrial shape, size, and function. Mitochondrial dynamics allows for the exchange of components, such as lipids and proteins, and the removal of damaged or defective mitochondria, maintaining the overall function and quality of mitochondria^1–5^.

Mitochondrial dynamics are controlled by dynamin-related GTPases: mitochondrial fission requires Dynamin Related Protein 1 (DRP1), while mitochondrial fusion depends on mitofusins (MFN1 and MFN2) for the fusion of the outer membrane and OPA1 for the fusion of the inner membrane. In addition to this core machinery, mitochondrial fission requires specific contacts with the endoplasmic reticulum (ER), which marks fission sites^6^. These ER-mitochondria contact sites then work as platforms to recruit the fission machinery. One key event driving mitochondrial fission is the activation of the ER-anchored formin protein Inverted Formin 2 (INF2), resulting in actin polymerization at ER-mitochondria contacts^7–9^. These actin filaments have been proposed to further promote ER-mitochondria contacts in a Myo19-dependent manner ^10^ and are used by non-muscle myosin II to constrict the mitochondrion^11, 12^. This then allows the recruitment, oligomerization, and activation of DRP1 and GTPase-dependent scission of the mitochondrion. Consistent with this model, inhibiting actin polymerization, myosin II activity, or INF2 decreases DRP1 oligomerization on mitochondria, suggesting that actin is involved in the recruitment and activation of DRP1 during mitochondrial fission^13–16^.

The role of ER and actin in mitochondrial fusion is less well understood. Recent studies have demonstrated that the ER is present not only at mitochondrial fission sites but also at fusion sites^17^ where it was suggested to accelerate the fusion process^18^. Consistent with this, loss of ER-mitochondria contact sites decreases the number of fusion and fission events^19^. Nevertheless, the exact role of the ER in mitochondrial fusion remains to be elucidated. One possibility is that it acts as a platform to recruit the fusion machinery, similarly to what occurs during fission^6, 7, 17^.

Previous research supports important roles for mitochondria-associated actin (“mito-actin”) aside from its role in mitochondrial fission. This includes mitochondria quality control^20^ and the regulation of mitochondrial transport^21–24^. Similarly, actin plays an important role in recruiting and activating dynamin family GTPases to various cell compartments^25, 26^, and has been shown to act in concert with dynamin to promote cell-to-cell fusion^27^. As the ER likely acts as a platform to recruit the machinery for both mitochondrial fission and fusion, and ER-anchored INF2 and actin play crucial roles in GTPase-mediated mitochondrial fission, we hypothesize that mito- and ER-associated actin (“ER-actin”) is important for GTPase-mediated mitochondrial fission and fusion.

Here, we uncover a previously unappreciated role for actin in the regulation of mitochondrial fusion, and provide new evidence for a role for both mito- and ER-actin in fission and fusion. Using live cell timelapse imaging of fluorescent protein-tagged mitochondria- and ER-targeted actin chromobodies, we demonstrate that actin marks the future sites of mitochondrial fission and fusion. Importantly, we show that disrupting actin filaments specifically on mitochondria or the ER disrupts both mitochondrial fission and fusion. In addition, inhibiting the actin nucleation protein complex Arp2/3 or ER-anchored formin protein INF2 differentially impacts the balance between mitochondrial fission and fusion. Our work also reveals two mechanistically distinct types of fusion events (tip-to-tip vs. tip-to-side) that can be distinguished based on their requirement for mitochondrial actin. We show that mito-actin is associated with the immobilized “receiving” mitochondrion during tip-to-side fusion, marking the fusion sites prior to the recruitment of the dynamin protein MFN2. Together, our data reveals a key organizational and regulatory role for mito- and ER-actin in both mitochondrial fission and fusion.

### Actin accumulates at sites of mitochondrial fusion

A classic way to assess mitochondrial fusion events is to measure the transfer of photo-activable GFP (PA-GFP) at sites of contact between two mitochondria. In primary human fibroblasts, 94% of contacts lasting more than 100 sec (114/121 fusion events in 12 cells) resulted in PA-GFP transfer between the two mitochondria (Supp. Figure 1, Video 1). To optimize the imaging setup with minimal phototoxicity, we used this validated 100 second criteria for the following experiments assessing mitochondrial actin.

To monitor the presence of actin at the site of mitochondrial fusion and fission, we used GFP-tagged actin nanobodies (termed Actin Chromobodies (AC)) targeted to the outer mitochondrial membrane with the c-terminal transmembrane domain of Fis1 (AC-mito)^8^. AC-mito moves freely within the mitochondrial membrane giving a diffused signal. However, when the AC probe binds to actin filaments within ∼10 nanometers of the mitochondrial outer membrane, it becomes immobilised, resulting in a higher intensity signal at the actin-associated site. We have previously used these probes to monitor the dynamics of actin accumulation at mitochondrial fission sites^8^. We transfected primary human fibroblasts with AC-mito and a control mCherry construct targeted to the mitochondrial outer membrane with the same Fis1 transmembrane domain as AC-mito (mCherry-mito) to label mitochondria and imaged them live with confocal microscopy. We first confirmed that AC-mito expression did not disrupt the balance between fusion and fission (although it did somewhat decrease overall mitochondrial dynamics) (Figure 1A). We then analysed the presence of the AC probe at mitochondrial fission and fusion sites. Consistent with previous studies, we observed AC-mito accumulation at fission sites (Figure 1B, total fission events/cells in Figure 1A; Video 2, stills in Supp. Figure 2A), validating the approach. Importantly, we also observed AC-mito accumulation at fusion sites in most fusion events (Figure 1B-C, total fusion events/cells in 1A; video 3). AC-mito was present at sites of mitochondrial fusion at a much higher frequency than what would be predicted by chance based on the area of mitochondria covered by AC-mito (Figure 1D), indicating that this is not a statistically random event^6, 8^. To confirm AC-mito accumulation at fusion and fission sites, we measured its specific enrichment relative to adjacent mitochondria. We observed a 2-fold increase in enrichment of AC-mito signal at fusion (Figure 1E) and fission (Figure 1F) events, which were absent when measured using the mCherry-mito signal, the latter consistently spreading evenly across mitochondrial surfaces (Figure 1E-F).

**Figure 1.**
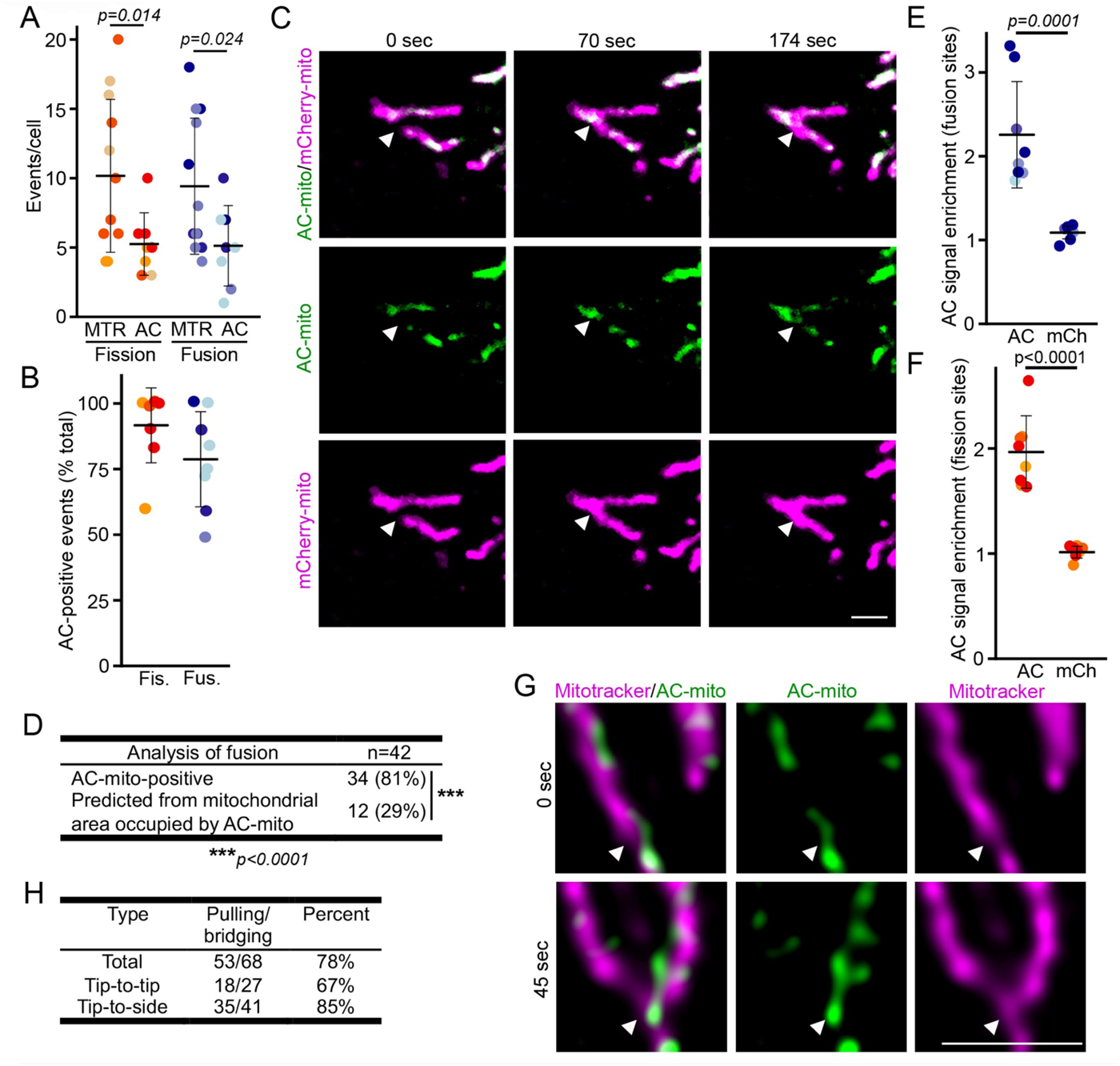
Actin marks the site of mitochondrial fusion. (A) Quantification of fission and fusion events in cells in which mitochondria were stained with Mitotracker (MTR) and cells transfected with mCherry-mito and AC-mito (AC). Each point represents an individual cell, with at least 10 cells quantified in 4 independent experiments. Bars show the average ±SD. (B-C) AC-positive fission and fusion events. Cells were transfected with mCherry-mito (mitochondria, magenta) and AC-mito (actin, green). Quantification of fission (Fis.) and fusion (Fus.) events positive for AC-mito. Each point represents an individual cell, with 10 cells quantified in 4 independent experiments (same cells as in (A) AC). Bars show the average ± SD. Two-sided t-test. Representative fusion event showing the enrichment of AC-mito at the fusion site (arrowhead) is shown in (C). Scale bar 2µm. (D) Number of AC-mito-positive fusion events compared to the expected number AC-mito-positive fusion events predicted from the mitochondrial area covered AC-mito). Fisher’s exact test. (E-F) AC-mito signal enrichment at fusion (E) and fission (F) sites in cells transfected as in B. Signal intensity at the event site relative to an adjacent site on the mitochondrial network was quantified for AC-mito (AC) and mCherry-mito (mCh). Each point represents an individual cell, with 8 cells quantified in 4 independent experiments. Bars show the average ±SD. Two-sided t-test. (G-H) SIM imaging of U2OS transfected with AC-mito and stained with mitotracker. (G) Representative image showing AC-mito (Green) and mitochondria (Mitotracker, Magenta). Scale bar 1 µm. The arrowheads point to the fusion site. (H) Quantification of the number of fusion events where actin bridges the two fusing mitochondria as in the bottom image in (G). The total number of fusion events from 5 cells is shown. The same data was also separated into tip-to-tip and tip-to-side fusion events (see Figure 2).

We then used lattice-structured illumination microscopy (lattice-SIM) on U2OS cells transfected with AC-mito and labelled with mitotracker to better assess the topology of actin accumulation at fusion sites. In these videos (Video 4), we observed AC-mito (green) labelling the future fusion site, then bridgingthe space between the two fusing mitochondria (mitotracker, magenta; Figure 1G). This pulling/bridging behavior was observed in the vast majority of fusion events (Figure 1H, total). In contrast, actin at fission sites was associated with pinching and separation of the two mitochondria (Video 5), consistent with previous reports ^6^. Overall, our data demonstrates that AC-mito is enriched at mitochondrial fusion sites, indicating the presence of mito-actin at these sites, and suggests that it could play a structural role in bridging the two fusing mitochondria.

Mitochondrial fusion occurs in two distinct ways: tip-to-tip fusion, where the end of one mitochondrion fuses with the end of another, and tip-to-side fusion, where the end of a mitochondrion fuses at the side of another mitochondrion^19^ (Figure 2A). Notably, our data revealed that 75% of fusion events involved tip-to-side fusion (Figure 2B). Interestingly, actin was much more likely to be present at tip-to-side events (88%) than end-to end events (50%) (Fig. 2C; Video 6 for tip-to-tip fusion, stills in Supp. Figure 2B), suggesting that actin predominantly favors tip-to-side fusion. Similarly, the pulling/bridging behaviour observed occurred most frequently in tip-to-side fusion events, but to a lesser extent in tip-to-tip fusion events (Figure 1H). We also observed that in most fusion events, one mitochondrion remains immobile while the other is mobile (Figure 2D, “Still”-immobile). This was however not the case for tip-to-tip fusions, where both mitochondria were usually mobile (Figure 2E).

**Figure 2.**
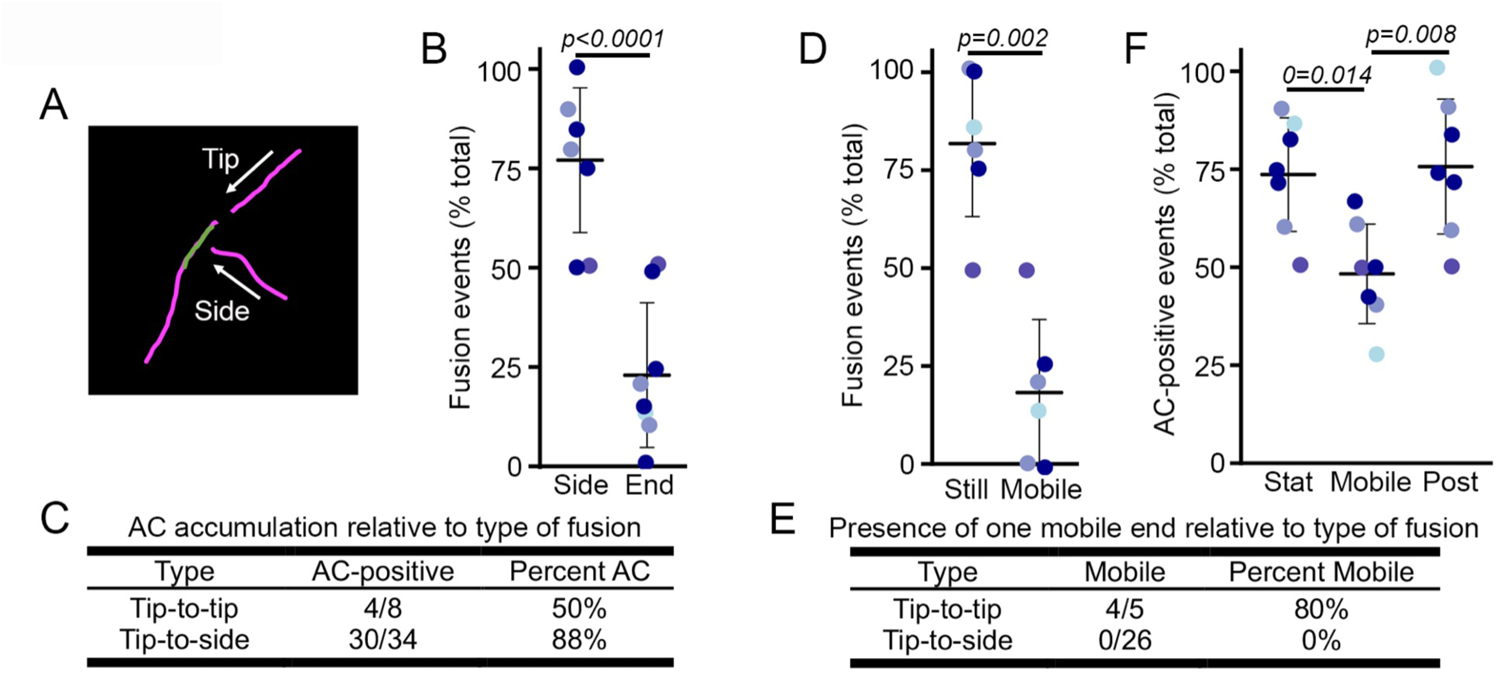
Actin is mainly associated with Tip-to-side mitochondrial fusion. (A) Schematic representation of Tip-to-tip and Side-to-End fusion events. (B-C) Quantification of fusion events as end to-side (Side) or tip-to-tip (End)(B), and their corresponding association with AC-mito (C) in cells from Figure 1B-C (transfected with mCherry-mito and AC-mito). Each point represents an individual cell, with 10 cells (E) quantified in 4 independent experiments. Bars show the average ± SD. Two-sided t-test. Total events from the quantified cells are shown in (C). (D-E) Quantification of fusion events containing one stationary mitochondrion (Still) or two mobile mitochondria (Mobile) (D) and their corresponding association with AC-mito (E). Each point represents an individual cell with 8 cells quantified in 4 independent experiments. Bars show the average ± SD. Two-sided t-test. Total events from the quantified cells are shown in (E). (F) Quantification of AC-mito signal associated with each of the fusing mitochondria (the stationary mitochondrion (Stat) and the mobile mitochondrion (Mobile)) as well as with the fusion site one frame post-fusion (Post). Each point represents an individual cell, with 9 cells quantified in 4 independent experiments. Bars show the average ± SD. One-way ANOVA.

Our subsequent objective was to identify which of the two fusing mitochondria recruited actin. Since most fusion events involved one immobile mitochondrion, we selected this subset for further analysis. We referred to the immobile mitochondrion as the “stationary mitochondrion” and the mobile mitochondrion as the “mobile mitochondrion”. Consistent with the lattice-SIM data showing AC-mito at the future fusion sites (Figure 1G), we observed that actin is present to a greater extent on the immobilized stationary mitochondrion than on the mobile mitochondrion, and that the AC-mito signal persists until the end of the fusion event (Figure 2F). Together, our findings indicate that mitochondria-associated actin acts as a marker for the sites of tip-to-side mitochondrial fusion.

### Actin-associated mitochondrial fusion sites recruit the endoplasmic reticulum (ER)

During mitochondrial fission, the ER acts as a platform to recruit actin-regulatory proteins (INF2) and actin-binding motor proteins (non-muscle myosin II) leading to mitochondrial constriction and DRP1-mediated scission of the mitochondrion^6, 7, 11, 13, 15, 28^. Accordingly, to determine if ER is providing a platform for fusion sites, we transfected primary human fibroblasts with mCherry targeted to the ER using the c-terminal domain of cytochrome b5 (mCherry-ER) and EGFP targeted to the mitochondrial matrix (mito-GFP), and imaged them live with confocal microscopy. Consistent with previous reports, most fusion and fission events were associated with the presence of ER (Figure 3A, total events in 3B, Videos 7 (fusion) and 8 (fission), stills in Supp. Figure 3A (fusion) and 3B (fission))^10, 17–19^. We then tested whether ER-actin also marked mitochondrial fusion sites. To specifically label ER-actin, we used our previously reported variation of the AC probe targeted to the ER using the c-terminal domain of cytochrome b5 (AC-ER)^8^. We simultaneously transfected cells with AC-ER and mCherry-mito and assessed the recruitment of AC-ER at fusion (Figure 3C, Video 9) and fission sites (Supp. Figure 4, Video 10). Similar to AC-mito, most fusion and fission events were positive for AC-ER (Figure 3D, total events in 3E). The enrichment of AC-ER was confirmed at fusion and fission sites (Figure 3F) as for AC-mito. Like AC-mito, most stationary mitochondria were AC-ER-positive, while mobile mitochondria were less likely to show AC-ER signal (Figure 3G). Together, our results indicate that both mito- and ER-actin are present at mitochondrial fusion sites.

**Figure 3.**
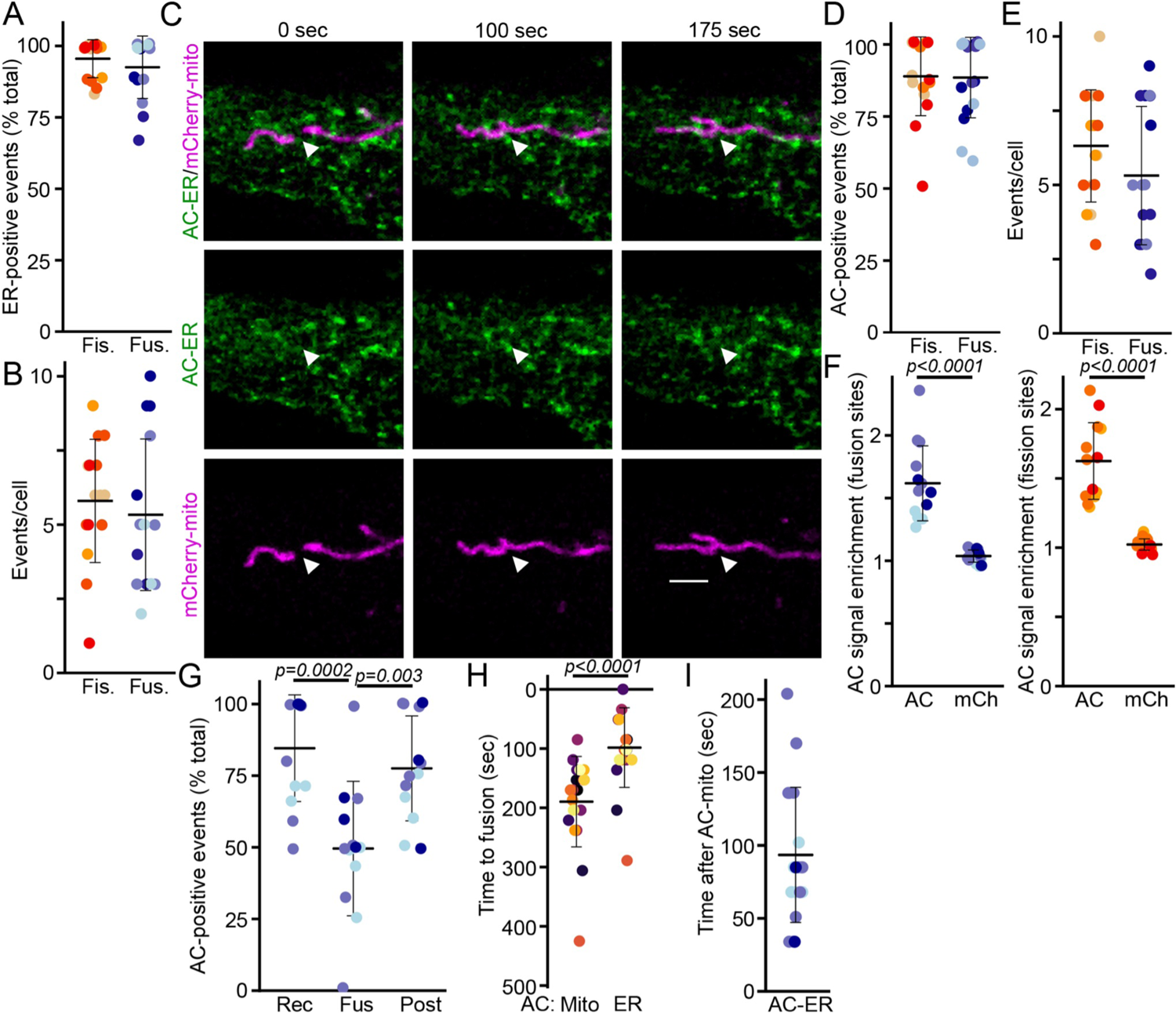
ER-associated actin is present at the site of mitochondrial fusion. (A-B) Quantification of fission (Fis.) and fusion (Fus.) events associated with ER. Cells were transfected with mCherry-ER and mito-GFP. Each point represents an individual cell, with 15 cells quantified in 4 independent experiments. (A) ER-Positive events (%), (B) Total events per cell. Bars show the average ± SD. (C) Representative fusion event showing the enrichment of AC-ER at the fusion site (arrowhead). Cells were transfected with mCherry-mito (mitochondria, magenta) and AC-ER (actin, green). Scale bar 2 µm. (D-E) Quantification of fission (Fis.) and fusion (Fus.) events positive for AC-mito. Each point represents an individual cell, with 13 cells quantified in 3 independent experiments. (D) ER-Positive events (%), (E) Total events per cell. (F) AC-ER signal enrichment at fusion (Left) and fission (Right) sites in cells transfected as in C. Signal intensity at the event site relative to an adjacent site on the ER network was quantified for AC-ER (AC) and mCherry-ER (mCh). Each point represents an individual cell, with 13 cells quantified in 3 independent experiments. Bars show the average ± SD. two-sided t-test. (G) Quantification of AC-ER signal associated with each of the fusing mitochondria (the stationary mitochondrion (Rec) and the mobile mitochondrion (Fus)) as well as with the fusion site one frame post-fusion (Post). Each point represents an individual cell, with 13 cells quantified in 3 independent experiments. Bars show the average ± SD. One-way ANOVA. (H-I) Kinetics of recruitment of AC-mito and AC-ER at the fusion site showing AC-ER recruitment relative to AC-mito (G) and recruitment of either marker relative to the time of fusion (H). Each point represents an individual event (20 events total in 4 cells within 3 independent experiments). two-sided t-test.

During fission events, mito-actin accumulates first at the fission site, followed by ER-actin ^8^. We similarly sought to determine the temporal relationship between the recruitment of AC-ER and AC-mito at fusion sites. To determine when mito- and ER-actin filaments are recruited to mitochondrial fusion sites, cells were co-transfected with AC-mito, a halo-tagged version of AC-ER, and mCherry-mito. We then measured ER- and mito-actin recruitment relative to fusion events. Similar to what was observed for mitochondrial fission, we found that AC-mito was recruited first to the future fusion site, followed by AC-ER (Figure 3H, fusion occurs at timepoint 0; (Supp. Figure 5, Video 11). Specifically, AC-ER was recruited at the fusion site at an average of 100 seconds after AC-mito (Figure 3I), suggesting that recruitment of actin on mitochondria, not the ER, is the primary event for both mitochondrial fission and fusion events.

### Mitochondria-associated actin is required for fusion

As our results show that actin marks mitochondrial fusion sites, we then asked whether it is required for fusion. Actin depolymerizing drugs impact many subcellular processes and signaling pathways, making it difficult to distinguish primary from secondary effects. Thus, the ability to selectively remove actin from subcellular compartments of interest would be enormously valuable for the cell biology research community. To address this gap and selectively remove actin from mitochondria, we modified a previously described <u>D</u>isassembly-promoting, <u>e</u>ncodable <u>Act</u>in tool (DeAct), a ∼120 amino acid domain of the actin-regulating protein Gelsolin (Gelsolin Segment 1 - aa 53-176 – GS1) that disrupts actin filaments ^29^. To specifically target GS1 to mitochondria or the ER, we fused it to the c-terminal transmembrane domain of Fis1 (DeAct-mito) or cytochrome b5 (DeAct-ER).

After confirming the proper localisation of the constructs (Supp. Figure 6A-B), we first determined if organelle-targeted DeAct prevented actin accumulation on mitochondria by co-transfecting DeAct-mScarlet (mito or ER) with AC-mito or AC-ER. Consistent with the ability of DeAct to locally disrupt actin filaments, DeAct-mito significantly decreased AC-Mito signal accumulation (Figure 4A-B), while AC-ER foci showed a steep decline when transfected with DeAct-ER (Figure 4C, Supp. Figure 7). We also tested whether the organelle-targeted DeActs specifically affected their target organelle. DeAct-mito did not significantly affect AC-ER signal, highlighting the selectivity of the probe (Figure 4A-C). In contrast, DeAct-ER was also efficient in preventing actin accumulation on mitochondria (Figure 4A-B), likely as a result of the ER making numerous contacts with mitochondria. Overall, both constructs remained selective for the relatively small organelle-bound pools of actin, as no obvious alterations were found in our phalloidin staining (Supp. Figure 6C).

**Figure 4.**
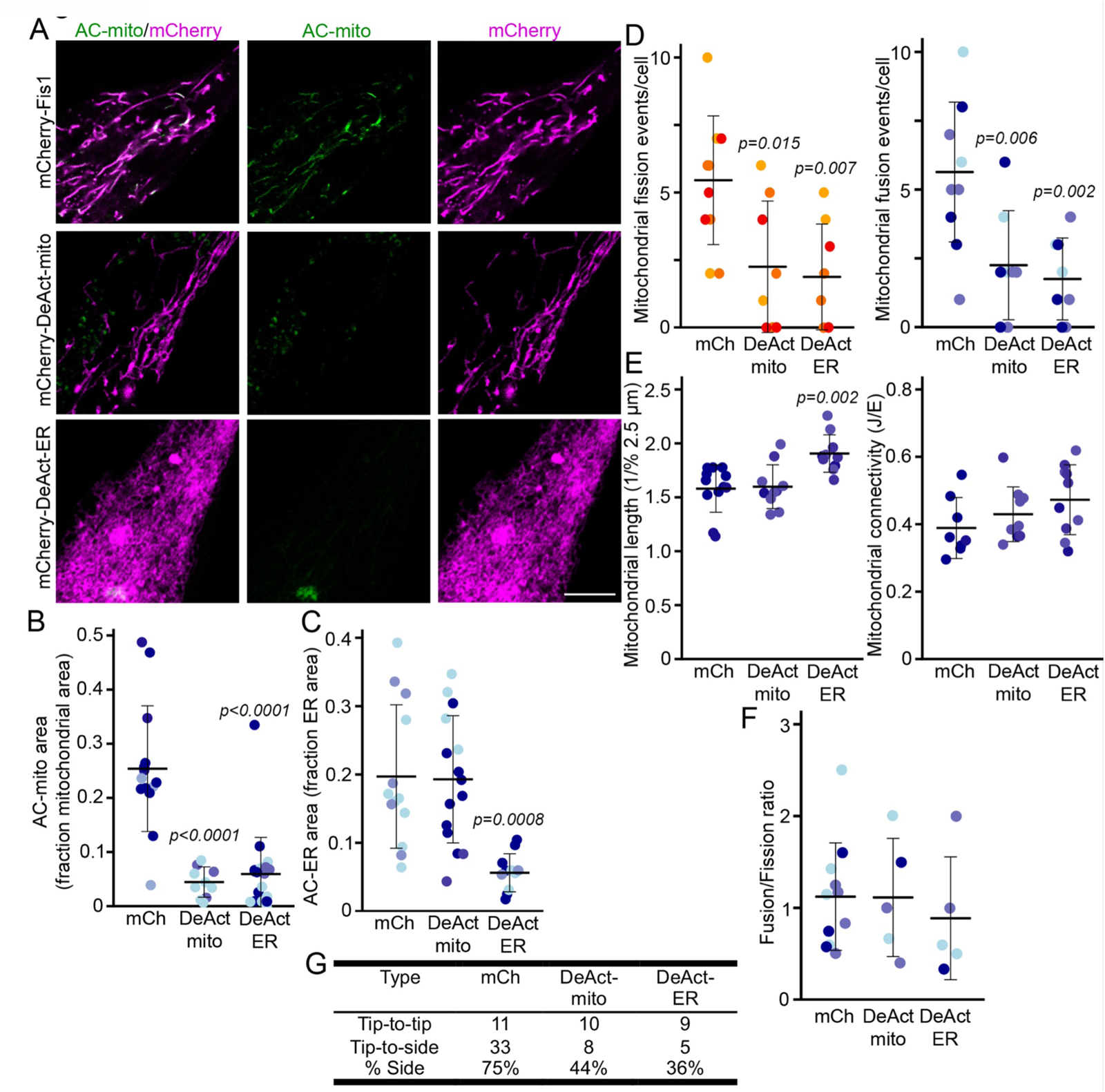
Mitochondrial actin is required for both fission and fusion. (A) Representative images showing the loss of AC-mito signal (green) in cells transfected with DeAct-mito or DeAct-ER (magenta) and AC-mito. (B-C) Quantification of AC-mito (B) and AC-ER (C) signal accumulation in cells transfected as in (A). Each point represents an individual cell, with at least 8 cells quantified in 3 independent experiments for each condition. Bars show the average ± SD. One-way ANOVA. (D) Quantification of the number of fission (Left) and fusion (Right) events in cells transfected with the indicated DeAct probe, AC-mito and mCherry-Fis1 (to label mitochondria). Each point represents an individual cell, with 11 (mCh), 8 (DeAct-mito) and 9 (DeAct-ER) cells quantified in 3 independent experiments. Bars show the average ± SD. One-way ANOVA. (E) Quantification of mitochondrial length (Left) and connectivity (Right) in cells from (D). Each point represents an individual cell. Bars show the average ± SD. One-way ANOVA. (F) fusion/fission ratio quantified from the data in (D). (G) Quantification of fusion events as tip-to-side (Side) or tip-to-tip (End). The total number of events for 8 cells in 3 experiments is shown for each condition.

Having validated the DeActs, we then evaluated their effect on mitochondrial fusion and fission rates. Consistent with the proposed role for ER-associated actin in mitochondrial fission, DeAct-ER significantly decreased mitochondrial fission rates (Figure 4D). DeAct-mito also inhibited fission (Figure 4D), suggesting that both ER and mitochondria-associated actin play a role in fission. Interestingly, DeAct-ER, but not DeAct-mito, led to elongated mitochondria (Figure 4E). Importantly, mitochondrial fusion was also significantly decreased by both DeActs (Figure 4D). As DeAct-mito did not impact ER-actin (Figure 4C), this indicates that mitochondria-associated actin is required for mitochondrial fusion. We then determined whether fusion or fission was more affected by DeActs by taking the ratio of fusion over fission for each condition. There was no significant difference in the ratio for either DeAct compared to control cells (mCherry-mito) (∼1), suggesting that DeActs blocked both processes (Figure 4F). These data therefore demonstrate an essential role for actin in the regulation of not only mitochondrial fission but also mitochondrial fusion.

Given the majority of fusion events that occurred in the presence of actin were of the tip-to-side pattern (Figure 2A-C), we asked whether one type of fusion event was more affected by DeActs. Consistent with tip-to-side events being mostly associated with actin, these were significantly affected by DeActs (Figure 4G). In contrast, tip-to-tip events persisted in DeAct-mito and DeAct-ER transfected cells (Figure 4G), suggesting that the residual fusion observed with DeActs is largely the consequence of actin-independent tip-to-tip fusion events.

### Arp2/3-dependent actin polymerization is required for mitochondrial fusion

Having demonstrated that actin is required for mitochondrial fusion, we next defined the role of actin-polymerizing proteins in this process. Most actin polymerization is regulated by one of two families of actin-regulatory proteins: Arp2/3 is usually responsible for the formation of branched actin filaments and actin patches, while formins usually promote the formation of parallel actin filaments.

To study the role of Arp2/3 in mitochondrial fusion, we selectively inhibited this nucleation complex using CK-666^30^. Firstly, there was a notable decrease in AC-mito accumulation following 60 min CK-666 treatment (Figure 5A-B), supporting a role for Arp2/3 in the generation of mito-actin. The CK-666-dependent loss of mito-actin was also associated with a significant decrease in the number of mitochondrial fusion events (Figure 5C). In contrast, CK-666 did not significantly affect mitochondrial fission (Figure 5C), resulting in a decrease in the fusion/fission ratio (Figure 5D). As the loss of mitochondrial actin due to the expression of DeAct-mito selectively inhibited tip-to-side fusion (Figure 4G), we also examined the effect of CK-666 on tip-to-tip and tip-to-side fusion. Consistent with the DeAct data, CK-666 selectively prevented tip-to-side fusion (Figure 5E). As tip-to-side fusion events increase the number of junctions within mitochondrial networks, they promote mitochondrial connectivity. Consistent with this, while CK-666 did not alter mitochondrial length, it significantly decreased mitochondrial connectivity (Figure 5F).

**Figure 5.**
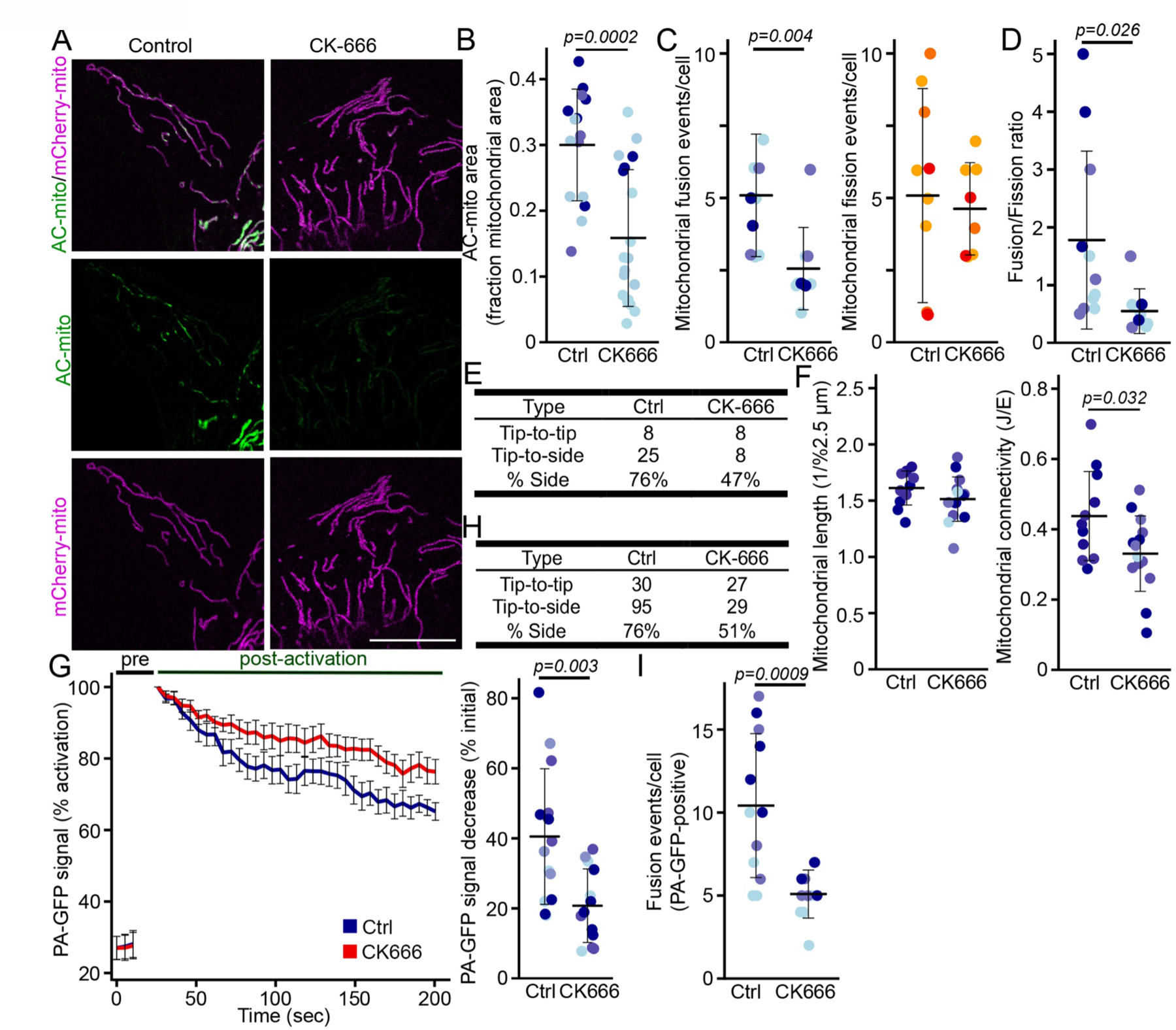
The Arp2/3 complex regulates mitochondrial fusion. (A) Representative images showing the loss of AC-mito signal in primary fibroblasts transfected with AC-mito (green) and mCherry-Fis1 (magenta), and treated with the Arp2 inhibitor CK666. Scale bar 10 µm (B) Quantification of AC-mito signal accumulation in cells transfected as in (A). Each point represents an individual cell, with 15 cells quantified per condition in 3 independent experiments. Bars show the average ± SD. Two-sided t-test. (C) Quantification of the number of fusion (Left, blue) and fission (Right, orange) events, as well as the fusion/fission ratio (D) in cells transfected as in (A) and treated as indicated. Each point represents an individual cell, with 12 (control) and 9 (CK666) cells quantified in 3 independent experiments. Bars show the average ± SD. Two-sided t-test. (E) Quantification of fusion events as tip-to-side (Side) or tip-to-tip (End). The total number of events for 9 cells in 3 experiments is shown for each condition. (F) Quantification of mitochondrial length (Left) and connectivity (Right) in cells from (C). Each point represents an individual cell. Bars show the average ± SD. One-way ANOVA. (G-I) Mitochondrial fusion assay. Human primary fibroblasts were transfected with photoactivatable-GFP (PA-GFP), treated as indicated and imaged before (pre) and after activation with the 405 nm laser. Fluorescence traces (G, Left) and the quantification of the loss of fluorescence at 3 min relative to the initial time post-activation (G, Right) are shown. Each point (G, Right) represents an individual cell, with 14 ctrl and 16 CK666 cells quantified in 4 independent experiments. Bars show the average ± SD. Two-sided t-test. Fusion was also directly quantified by measuring the events in which PA-GFP was transferred from one mitochondrion to another upon fusion (H, separation by fusion type; I, total fusion events). Each point represents an individual cell, with 12 ctrl and 11 CK666 cells quantified in 3 independent experiments. Bars show the average ± SD. Two-sided t-test.

To determine whether our observations were cell-specific, we also tested the effect of CK-666 in the breast cancer line MDA-MB-231. These cells showed a similar decrease in mitochondrial fusion following CK-666 treatment (Supp. Figure 8A). To further confirm our findings, we knocked down Arp2 in MDA-MB-231 cells (Supp. Figure 8C) and quantified mitochondrial fission and fusion. Consistent with the CK-666 data, mitochondrial fusion was significantly reduced in cells where Arp2 was knocked down (Supp. Figure 8D). In contrast, mitochondrial fission was not significantly affected (Supp. Figure 8D), leading to a decrease in the fusion/fission ratio (Supp. Figure 8E).

To further demonstrate that the Arp2/3 complex is required for mitochondrial fusion, we used the PA-GFP mitochondrial fusion assay (Figure 5G, Supp. Figure 9). Consistent with Arp2/3 inhibition preventing mitochondrial fusion, PA-GFP signal showed a lower rate of decrease in CK666-treated cells compared to control cells, leading to a significantly smaller decrease in GFP signal 3 minutes after activation (Figure 5G). We also confirmed the inhibitory effect of CK-666 on mitochondrial fusion by quantifying the transfer of photo-activated GFP from one mitochondrion to another during fusion and showing that CK-666 prevents PA-GFP transfer selectively for tip-to-side fusion events (Figure 5H-I). Altogether, our data indicate that Arp2/3-dependent actin polymerization plays a crucial role in regulating mitochondrial fusion.

### Formin inhibition predominantly affects mitochondrial fission

While our results demonstrate a role for Arp2/3 in mitochondrial fusion, formin proteins are also known to be important for actin dynamics and can both collaborate and compete with Arp2/3 in forming dynamic actin structures^31–33^. In addition, both Arp2/3 and formin proteins are known to be important for mitochondrial fission dynamics^34–36^. We thus asked whether formin-mediated actin polymerization could also play a role in mitochondrial fusion. For this, we first used the Formin FH2 Domain Inhibitor (SMIFH2) to inhibit formin-mediated elongation of actin filaments. As with Arp2/3 complex inhibition, formin inhibition resulted in a significant loss of AC-mito accumulation (Figure 6A-B). Consistent with the role of formins in mitochondrial fission, fission was also disrupted by SMIFH2 (Figure 6C). On the other hand, while SMIFH2 did reduce mitochondrial fusion, this effect was lower than for fission, leading to a significant increase in the fusion/fission ratio (Figure 6C-D). Consistent with this, SMIFH2 caused mitochondrial elongation but no change in mitochondrial connectivity (Figure 6E). Nevertheless, as for Arp2/3 inhibition, SMIFH2 selectively prevented tip-to-side fusion events (Figure 6F), consistent with actin being predominantly required for this type of fusion event.

**Figure 6.**
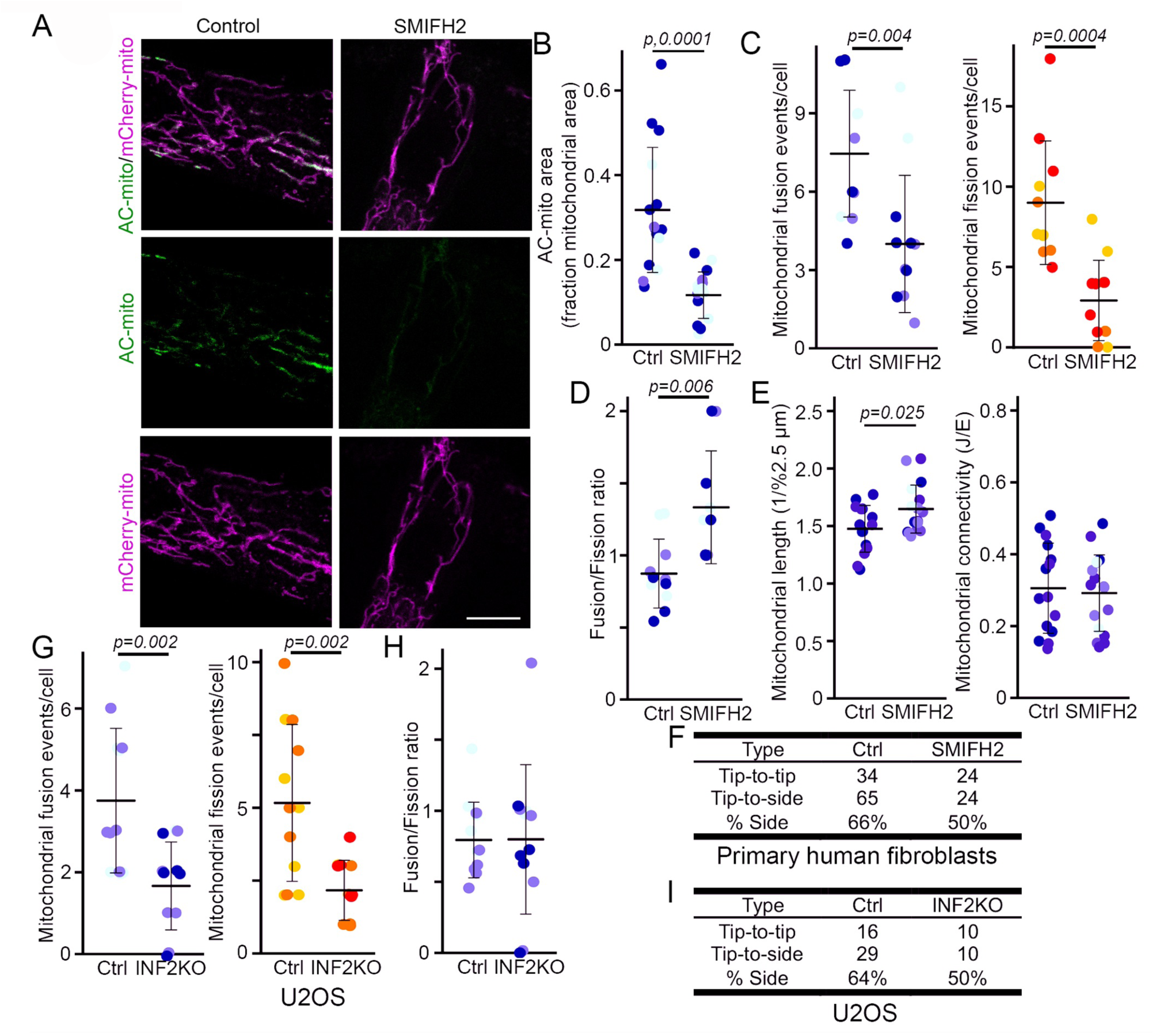
Formin-dependent regulation of mitochondrial fusion. (A) Representative images showing AC-mito signal in primary fibroblasts transfected with AC-mito (green) and mCherry-Fis1 (magenta), and treated with the formin inhibitor SMIFH2. Scale bar 10 µm (B) Quantification of AC-mito signal accumulation in cells transfected as in (A). Each point represents an individual cell, with 12 cells quantified in 3 independent experiments. Bars show the average ±SD. Two-sided t-test. (C-D) Quantification of the number of fusion (C, Left, blue) and fission (C, Right, orange) events, as well as the fusion/fission ratio (D) in cells transfected as in (A) and treated as indicated. Each point represents an individual cell, with 12 cells quantified in 3 independent experiments. Bars show the average ± SD. Two-sided t-test. (E) Quantification of mitochondrial length (Left) and connectivity (Right) in cells from (B). Each point represents an individual cell. Bars show the average ± SD. Two-sided t-test. (F) Quantification of fusion events as tip-to-side (Side) or tip-to-tip (End). The total number of events for 12 cells in 3 experiments is shown for each condition. (G-H) Quantification of the number of fusion (Left, blue) and fission (Right, orange) events, as well as the fusion/fission ratio (H) in U2OS cells in which INF2 has been deleted using CRISPR/Cas9. Each point represents an individual cell, with 12 cells quantified in 3 independent experiments. Bars show the average ± SD. Two-sided t-test. (I) Quantification of fusion events as tip-to-side (Side) or tip-to-tip (End). The total number of events for 12 cells in 3 experiments is shown for each condition.

The formin protein INF2 is an ER-anchored actin regulatory protein and is required for mitochondrial fission^7, 9^. We thus specifically examined the role of INF2 in fusion by measuring fission and fusion events in U2OS cells lacking INF2^9^. Consistent with our formin inhibitor data, INF2 deletion affected both fission and fusion (Figure 6G). However, contrary to the formin inhibitor, both processes were affected to a similar extent, resulting in a fusion/fission ratio that was similar between control and INF2 KO cells (Figure 6H). Importantly, as with our other manipulations of the actin cytoskeleton, INF2 deletion selectively affected tip-to-side fusion (Figure 6I). Altogether, our results indicate that actin is required for tip-to-side events which constitute the majority of fusion events within cells, and that both Arp2/3 and INF2 play a role in this process.

### Actin polymerisation on mitochondria precedes MFN2 recruitment to the fusion site

Mitochondrial fusion requires the action of MFN1/2 on the mitochondrial outer membrane and OPA1 on the mitochondrial inner membrane. Recently, both fission and fusion have been suggested to occur at hotspots where both fusion (MFN1/2) and fission (DRP1) GTPases coalesce at ER-mitochondria contact sites ^17^. To determine the relationship between these hotspots and mito-actin, we first determined the relationship between MFN2 recruitment and AC-mito signal at sites of mitochondrial fusion and fission. For this, we transfected primary fibroblasts with mCherry-MFN2 and AC-mito, stained mitochondria with mitotracker deep red, and determined the presence of each probe at fusion and fission sites (Supp. Figure 10, Video 12 - fission, Video 13 - fusion). Both MFN2 and AC-mito were present on the vast majority of fission and tip-to-side fusion events, consistent with the presence of fission/fusion hotspots (Figure 7A). During both fission and tip-to-side fusion, mito-actin marked fission/fusion sites 60-90 seconds prior to MFN2 recruitment (time to fission, Figure 7B-C; time relative to AC-mito, Figure 7D). This was however less clear for the tip-to-tip fusion events, which do not require mito-actin (Figure 7C-D).

**Figure 7.**
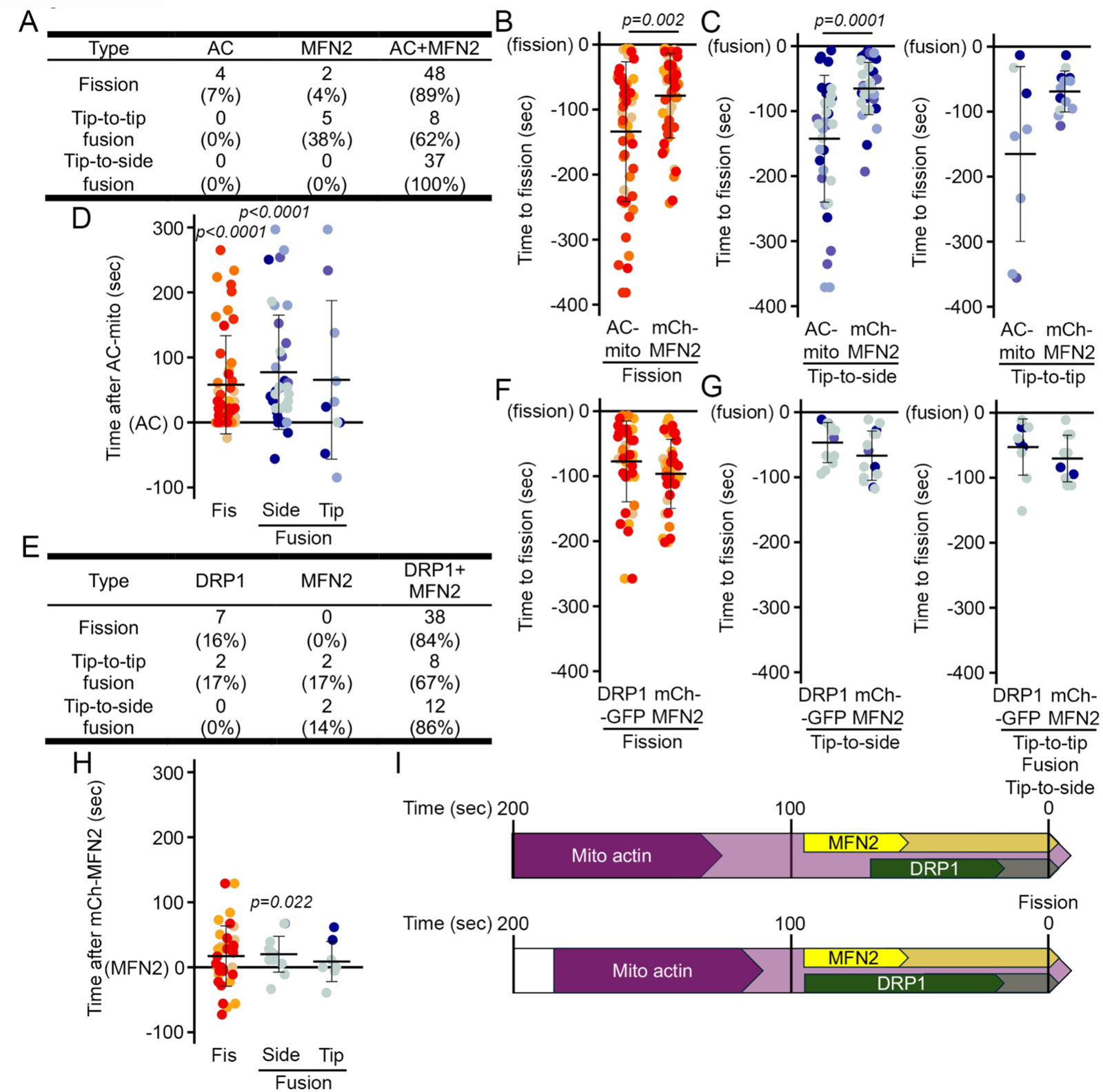
Mitochondrial actin precedes the recruitment of mitochondrial Dynamins at fusion and fission sites. (A-D) Temporal relationship between mitochondrial actin and the fusion dynamin MFN2. Cells were transfected with mCherry-MFN2 and AC-mito while mitochondria were visualised with Mitotracker Deep red. (A) Presence of AC-mito and mCherry-MFN2 at sites of mitochondrial fission, tip-to-side fusion and tip-to-tip fusion. (B-D) Kinetics of recruitment of AC-mito and mCherry-MFN2 at fission (B) and fusion (C) sites showing recruitment of either marker relative to the time of fusion, as well as mCherry-MFN2 recruitment relative to AC-mito (D). Each point represents an individual event in 4 cells within 3 independent experiments). two-sided t-test. (E-H) Temporal relationship between the fusion dynamin MFN2 and the fission Dynamin DRP1. Cells were transfected with mCherry-MFN2 and GFP-DRP1 while mitochondria were visualised with Mitotracker Deep red. (E) Presence of GFP-DRP1 and mCherry-MFN2 at sites of mitochondrial fission, tip-to-side fusion and tip-to-tip fusion. (F-H) Kinetics of recruitment of mCherry-MFN2 andGFP-DRP1 at fission (F) and fusion (G) sites showing recruitment of either marker relative to the time of fusion, as well as GFP-DRP1 recruitment relative to mCherry-MFN2 (D). Each point represents an individual event in 3 cells within 3 independent experiments). two-sided t-test. (I) Schematic representation of the timing of MFN2 and DRP1 recruitment to tip-to-side fusion (Top) and fission (Bottom) relative to actin recruitment (AC-mito-positive).

We then determined the relationship between MFN2 and DRP1 at these mito-actin hotspots. For this, we transfected primary fibroblasts with mCherry-MFN2 and GFP-DRP1, stained mitochondria with mitotracker deep red and determined the presence of each probe at fission and fusion sites (Supp. Figure 11, Video 14 - fission, Video 15 - fusion). Consistent with mito-actin marking fission/fusion hotspots, DRP1 and MFN2 were present at most fission and fusion sites, although the correlation was somewhat lower at actin-independent tip-to-tip fusion events (Figure 7E). MFN2 and DRP1 were recruited at similar times before fusion or fission (time to fission, Figure 7F-G). However, for tip-to-side fusion, DRP1 was generally recruited after MFN2 (time to fission, Figure 7F-G), although the time between the two events was relatively short (20 seconds, time relative to mCherry-MFN2, Figure 7H). These findings suggest mito-actin marks the sites of fission/fusion hotspots prior to the recruitment of ER and the fission/fusion GTPases (Figure 7I).

## Discussion

Actin plays a critical role in regulating organelle dynamics within cells, contributing to their movement, positioning, and organization^37–39^. Actin also plays an important role in mitochondrial fission, where it facilitates constriction of mitochondria and the recruitment and activation of DRP1 GTPase activity ^9, 15, 16, 40^. The recruitment of actin to fission sites was originally shown under fission-inducing conditions, but has recently been demonstrated under unstimulated conditions by the use of organelle-targeted actin chromobodies ^8^. It was also proposed that the requirement for actin filaments for mitochondrial fission is context-dependent, with only ∼50% of fission events coinciding with actin polymerization^41^. However, Schiavon et al, 2020 and this current study revealed a much higher incidence of actin filaments at fission sites, likely as a result of using the more sensitive organelle-targeted actin probes.

In this study, we provide strong evidence that ER- and mito-actin are required for both mitochondrial fission and fusion. Using mitochondria- and ER-targeted AC probes ^8^ we demonstrate that mito-actin is recruited to mitochondria during both fusion and fission under unstimulated conditions. The ER and ER-actin are also present at fusion sites, but they are recruited at pre-existing actin rich sites on mitochondria just before or during fusion. Similarly, both MFN2 and DRP1 were recruited to fusion and fission sites that were already marked with mito-actin, indicating that actin polymerisation acts as an important early step during both mitochondrial fission and fusion. In addition to this compelling temporality, we further demonstrated a causal role for actin using our novel genetically-encoded DeAct-ER/mito tools to selectively disrupt ER- and mito-actin. We found that selectively disrupting mito-actin blocked both mitochondrial fission and fusion, demonstrating the pivotal role of mito-actin in both processes. Together, our results reveal a heretofore entirely unknown role for actin in fusion.

The role of ER- and mito-actin during fission is thought to be relatively straightforward, as it is required to partially constrict the mitochondrion at the fission site and facilitate DRP1 recruitment^6^. In contrast, our imaging results indicate that, during fusion, mito-actin bridges the two fusing mitochondria, thus playing a potential structural role in the fusion process. In this context, actin could stabilize or hold the two mitochondria in place to enable fusion to occur. On the other hand, some myosins including Myo19 have been shown to regulate mitochondrial dynamics^42, 43^. Similarly, in yeast^21, 44^ and higher plants^22, 45^, actin filaments can interact with specific motor proteins to regulate short-distance mitochondrial movement, which could also serve to properly align the two fusing mitochondria. In this context, mito- and ER-actin, which are both recruited to fusion sites, may act at different steps in the recruitment of the fusion machinery and stabilization of the fusing mitochondria. In addition, recent studies have suggested that the actin cytoskeleton is crucial for maintaining the curvature of mitochondrial membranes^23^, which could also be important for the fusion process^46^.

In addition to a structural role, the fact that the formation of mitochondria-associated actin filaments is the initial step for the fusion process suggests that it could also participate in the recruitment or activation of the fusion machinery. This would allow the coordinated recruitment of the different cellular components required for fusion. As an example, the dynamic interaction between actin and dynamin promotes the bundling of Arp2/3-dependent actin filaments in a manner that is dependent on cycles of GTP hydrolysis by dynamin^27^. The actin-dependent promotion of the GTPase activity of the dynamin-family protein DRP1 is also relatively well-established in mitochondrial fission (https://doi.org/10.7554/eLife.11553). Together, these prior results along with the observed recruitment of MFN2 to mito-actin sites during mitochondrial fusion is consistent with a model wherein mitofusin dynamin GTPase proteins are recruited and activated by mito-actin at fusion sites.

Mitochondria-associated actin has previously been reported in a number of settings in addition to mitochondrial fission. For example, actin clouds assemble around depolarised mitochondria to isolate them for autophagy^20^, while mitochondrial actin comet tails promote mitochondrial redistribution during mitosis^36^. Distinct actin-polymerizing complexes are likely required depending on the cellular context. For example, two distinct populations of actin filaments have been suggested to regulate mitochondrial fission, the INF2-dependent pathway and a second, INF2-independent but Arp2/3-dependent pathway^9, 35^. These mitochondria-associated actin polymerization processes differ in the cellular stimuli resulting in fission^7, 9, 35^. On the other hand, some studies showed that both formins and Arp2/3 are involved in complex-mediated actin assembly around mitochondria^9, 34, 47^. Overall, multiple mechanisms activated on either mitochondria or ER-mitochondria contact sites likely cooperate to stimulate the formation of mitochondria-associated actin filaments. We propose that these filaments are used to organise downstream events, including fission and fusion. The specific outcome almost certainly depends on the spatial and temporal organisation of the machinery recruited downstream of mito-actin. For example, actin- and ER-dependent constriction of a mitochondrial tubule occurs at fission sites, while the actin patch present at fusion sites likely bridges or stabilizes the two fusing mitochondria.

This spatio-temporal regulation of fusion is also illustrated by the presence of two types of mitochondrial fusion events with different requirements for actin. Tip-to-side fusion, which constitute the majority of fusion events and lead to the formation of branched mitochondria, predominantly occurred in an actin-dependent fashion. On the other hand, tip-to-tip fusion events, which elongates mitochondria, were much less likely to show actin dependence. Importantly, disrupting actin polymerization significantly decreased the occurrence of tip-to-side fusion events, but not tip-to-tip events, demonstrating the selectivity of actin for tip-to-side fusion. This was also reflected in the selective changes observed in mitochondrial networks following Arp2/3 complex or formin inhibition, the first causing a loss of mitochondrial connectivity and the latter, increased mitochondrial length. Recent studies have demonstrated that the deletion of ABHD16A, an ER protein regulating the formation of fission and fusion nodes containing MFNs and DRP1 at ER-mitochondria contact sites, results in the loss of tip-to-side fusion ^19^. This suggests that actin could facilitate the formation of these nodes, therefore promoting tip-to-side fusion. Consistent with this, we observed the recruitment of both MFN2 and DRP1 to mito-actin patches. These observations may potentially serve as the basis for new insights into genetic and environmental conditions that result in altered branching in mitochondrial networks.

Some studies have noted the presence of elongated mitochondria upon inhibition of actin-polymerizing proteins. While this was seen as evidence for a loss of fission relative to fusion, our results suggest that tip-to-tip fusion could occur in the absence of actin, thus promoting mitochondrial elongation rather than branching. The dynamic nature of mitochondrial fission and fusion and the requirement for cells to maintain a balance between the two must also be considered. For example,the deletion of INF2 resulted in a reduction of both fusion and fission events. Under these conditions (steady state loss of INF2), it is plausible that the decreased fission rates led to a subsequent reduction in fusion events to maintain a proper mitochondrial network. Conversely, acute inhibition of Arp2/3 (one hour treatment) showed persistent fission events in cells, possibly due to INF2-dependent fission mechanisms^7, 9, 28, 35^. Whether or how cells adapt to chronic versus acute changes to fission/fusion machineries is an important question for future studies.

In summary, our findings reveal a new role for actin polymerization in organizing tip-to-side fusion events. The mechanism of actin promoting fusion remains yet to be elucidated, although we speculate it likely involves the stabilisation of the fusing mitochondria and the recruitment of the fusion machinery. Further research is needed to understand the mechanism and physiological implications of this discovery. Our study highlights the multifunctionality of actin in regulating organelle dynamics, emphasizing the need for continued tool development and investigation in this area.

## Material and Methods

### Cell culture

Primary Human Fibroblasts were purchased from the Coriell institute and cultured in Dulbecco’s modified Eagle’s medium (DMEM) containing 10% fetal bovine serum (FBS), supplemented with Penicillin/Streptomycin (100 IU/ml/100µL/mL). MDA-MB-231 cells (triple negative breast cancer) were purchased from ATCC and cultured in DMEM supplemented with 10% fetal bovine serum. These cells were stably transfected with mitochondria-targeted GFP (CCOeGFP, addgene, pLVX-EF1a-CCO-IRES, #134861) using Metafectene (Biontex). Cell culture reagents were obtained from Wisent.

### Plasmids and Transient transfection of primary cells

Primary cells were trypsinized and centrifuged at 5000 rpm for 5 minutes. Cell pellets were suspended in 10µl of the Neon transfection buffer R (ThermoFisher Scientific, MPK1096). Cells were transiently transfected with mito-GFP (Cox8A-EGFP) (Addgene #134861), mito-PAGFP (Addgene, #23348), mCherry-MFN2 (Addgene #141156), GFP-DRP1 (Addgene #191942), mCherry-Fis1^8^, mCherry-Cytb5^8^, Halo-Fis1^8^ and all custom actin chromobody probes that we previously reported^8^ using the Neon transfection system (ThermoFisher Scientific, MPK5000) as described ^48^. For all experiments, 1 µg of total DNA concentrations were transfected per 10 ul of 10^6^ cells/ml (individually or combined for co-transfection). Live cell imaging or immunofluorescence (IF) was performed 24 hours after transfection. However, for GFP-DRP1 and mCherry-MFN2, imaging was restricted to within 8 hours post-transfection due to significant adverse effects on mitochondrial networks and motility observed at longer time points. DeAct-ER and DeAct-Mito constructs where generated utilizing the “Gelsolin Segment 1 (GS1)” sequence described in Harterink et al^29^. CMV-DeAct-GS1 was a gift from Brad Zuchero (Addgene plasmid # 89445). The GS1 sequence was cloned into the pSIN vector under a CMV promoter, N-terminal to the mScarlet fluorescent protein sequence. Either a Cytb5ER or Fis1 localization sequence was fused to the mScarlet C-terminus to target expression to the ER (DeAct-ER) or mitochondrial outer membrane (DeAct-Mito), respectively. The targeting sequences for DeAct-ER/Cytb5ER (IDSSSSWWTNWVIPAISAVAVALMYRLYMAED) and DeAct-mito/Fis1 (IQKETLKGVVVAGGVLAGAVAVASFFLRNKRR) were the same as those previously described in Schiavon et al.^8^ to target AC-ER and AC-Mito to the ER or mitochondrial outer membrane.

### siRNA treatment

MDA-MB-231 were seeded onto 6 well dishes to reach 30–40% density in 24 hours. Then the cells were transfected with 10 nM of Arp2 siRNA (Dharmacon reagents, ACTR2 Gene 10097) and control siRNA (Thermo Fisher Scientific, Silencer Select, 4390843) using siLenFect lipid reagent (Bio-Rad,1703361). After 24 hours, the cells were imaged live or collected for western blotting.

### Drug treatments

Cells were incubated with 10 µM CK-666 (CAS 442633-00-3 – Calbiochem) or Formin FH2 Domain Inhibitor, SMIFH2 (CAS 340316-62-3 – Calbiochem) for 60 minutes to inhibit Arp2/3 complex and formins, respectively.

### Immunofluorescence

Transfected cells were seeded onto glass coverslips (Fisherbrand, 1254580) and allowed to adhere overnight. Cells were fixed with 4% paraformaldehyde for 15 minutes at room temperature (RT). Cells were then permeabilized with 0.2% Triton X-100 in PBS and blocked with 1% BSA / 0.1% Triton X-100 in PBS. The cells were then incubated with primary antibodies using following antibodies as per the experiment: TOM20 (Rb, Abcam, ab186735, 1:250), RTN4/NOGOA (Rb, Bio-Rad, AHP1799, 1:200), followed by with fluorescent tagged secondary antibodies (Jackson Immunoresearch, 1:500) and DAPI (Invitrogen, Thermo Fisher, D1306, 1:100).

### Live imaging and confocal microscopy

For Live cell microscopy imaging, cells were grown on glass bottom dishes in complete medium and stained for 30 min with 250 nM TMRM (Thermo fisher Scientific, T668) or 5 min with 100nM Mitotracker Deep Red (Thermo fisher scientific, M7512) where indicated. After staining, cells were washed 3 times with pre-warmed 1× phosphate buffered saline (PBS), and normal growth media was added prior to imaging. The plates were mounted onto a Leica TSC SP8 confocal microscope fitted with a 63×/1.40 oil objective using the optimal resolution for the wavelength (determined using the Leica software). Time-lapse images were acquired at a speed of (0.05-0.125 frames/s) for 10 minutes. For the MFN2/DRP1 transfection experiment, cells were imaged with Plan-Apochromat 63x/1.40 Oil DIC M27 objective on an inverted Zeiss 980 LSM Airyscan confocal microscope. Time-lapses were acquired at a speed of 5-11.5sec frames/s for 10min. Following acquisition, images were Airyscan processed using auto-filter 2D-SR settings in Zen Blue.

For the INF2 experiments, Wild-type and INF2 KO U2OS cells^9^ were maintained in DMEM supplemented with 10% FBS at 37C with 5% CO2. Cells were plated on 8-well no. 1.5 imaging chambers (Cellvis) coated with 10ug/mL fibronectin. Cells were stained with 50nM MitoTracker Deep Red for 30min, washed with PBS, then imaged in FluoroBright medium (Thermo Fisher) supplemented with 10% FBS. Cells were imaged with a C-Apochromat x40/1.2 NA W Korr FCS M27 objective on an inverted Zeiss 980 LSM Airyscan confocal microscope with the environmental control system supplying 37C, 5% CO2, and humidity for live-cell imaging. MitoTracker Deep Red was imaged with a 647-nm laser line at ∼250nW laser power at optimal resolution (determined by Zen Black software). Time-lapses were acquired at a speed of 0.152-0.357 frames/s for 5min. Following acquisition, images were Airyscan processed using auto-filter 2D-SR settings in Zen Blue.

For the lattice-SIM imaging, a ZEISS Elyra 7 using a 63x/1.4 oil DIC M27 objective (RI: 1.518) was employed. Images were processed using the SIM2 algorithm on 15 phase images. Two PCO Edge 4.2 cameras independently and simultaneously collected data from the 488 nm and 642 nm excitation channels, with a Dual Camera Beam Splitter (SBS LP 560) utilized for channel separation.

### Image processing and analysis

All image processing and analyses were done in ImageJ/Fiji. The images shown are from single focal planes unless stated otherwise. For mitochondrial area, the images were segmented in ImageJ using Filter/Median (1.0), then thresholding and adjusting the resulting image using Binary/Erode. Total mitochondria area was then measured using the Fiji Measure function.

### Tracking of Fusion and Fission events

Quantification of fusion and fission events was done manually. The images were tracked frame by frame to record movement of mitochondria that resulted in fusion or fission. Stable separation of the mitochondrial marker was counted as a fission event. Fusion events were defined as follows: two fusing mitochondria must retain their fused state for the following 10 frames (or at least 100 sec). If there was a division at the fusion site within 5 frames, such an event is discarded. Also, the mean signal intensity of the mitochondrial marker at the site of fusion and a region elsewhere on mitochondria (basal) was recorded at the end of the 5 frames. If the ratio of signal intensity at the site of fusion and basal signal intensity was higher than 1, it was considered as an overlapping, not a fusion event. The presence of ER at the site of fusion or fission was also assessed manually, by looking for ER colocalizing with mitochondria at the site.

### Tracking of AC-mito and AC-ER at fusion sites

To identify when actin is accumulated on ER and mitochondria relative to a fusion event, we counted backwards from that fusion event to determine at which frame the AC probe accumulated on mitochondria and ER. Determining the time between the frames where AC-mito and AC-ER appear at the fusion site gives us the information on the time taken by Actin to get accumulated on the organelle.

### AC probe ‘accumulation’ calculation

Using Fiji, a square selection was drawn around a region with obvious AC probe accumulation (0.8 µm^2^ for mitochondria and ER). The mean pixel intensity of AC-GFP and mCherry channels was measured within the selection. Another square of equal dimensions was drawn in an adjacent, control area with mCherry signal and mean pixel intensity was measured. The mean pixel intensity in the accumulated region was then divided by the mean pixel intensity in the control region.

### AC Signal

The changes in AC probe signal was determined in cells transfected with DeActs or upon treatment with inhibitors for Arp2/3 complex and formins. The area of mitochondria or ER was determined using mCherry-mito or mCherry-ER (or the DeAct probes when present). The mask of the organelle signal was then applied to the AC signal and the AC signal thresholded to select the areas of AC accumulation. The area of the AC signal was then divided by the organelle area to determine the fraction of the organelle area covered by the AC signal. The imaging parameters (laser intensity and gain values) were kept identical for control and test conditions during image acquisition for every experiment.

### mtPA-GFP-based Mitochondrial Fusion Assay

The photoactivation-based fusion assay^49^ was conducted by expressing Mito-PAGFP (Addgene #23348) in primary human fibroblasts. The cells were co-stained with TMRM to monitor mitochondrial membrane potential. Photoactivation was performed in selected ROIs using the 405 nm laser with 100% power and 40 iterations. Upon activation of mito-PAGFP, a small portion of the mitochondrial network is photoactivated, and the spread or loss of the signal to the rest of the mitochondrial network was recorded in the ROIs. The signal of all ROIs with significant PAGFP activation within a cell was then averaged and normalized to the initial signal intensity post-activation.

### Mitochondrial length and connectivity

Mitochondrial length and connectivity was quantified using the Momito algorithm^50^. Mitochondrial networks were first segmented using the Tubeness plugin and default thresholding. Images were then processed using Momito. Connectivity was calculated using the ratio of junctions (SE3) over ends (SE1).

### Western Blot

Cells were lysed in 10 mM Tris-HCl, pH 7.4, 1mM EDTA, 150 mM NaCl, 1% Triton X-100, complemented with a protease inhibitor cocktail (Sigma-Aldrich) and phosphatase inhibitor (Sigma-Aldrich), kept on ice for 10 min and centrifuged at 13000rpm for 10 minutes. Protein supernatants were collected, and protein concentration was estimated by DC protein assay kit (BioRad). For SDS-PAGE, 30 µg of proteins were mixed with 1X Lammeli buffer containing β-mercaptoethanol, then subjected to SDS-PAGE, transferred to a nitrocellulose membrane and blotted with the indicated antibodies (Arp2 (Santacruz, (E-12): sc-166103, 1:1000), HRP-tagged Actin (1:10,000). Membranes were then incubated with a 1:5000 dilution of horseradish peroxidase-conjugated secondary antibodies (Jackson Immunoresearch) and visualized by enhanced chemiluminescence (Thermo Fisher scientific) using a Bio-Rad imaging system.

## Data analysis and statistics

All graphs and statistical analysis were done using R. Immunofluorescence data were quantified and images representative of at least three independent experiments shown (exact n are in the quantification figures). Data is represented as average ± SD as specified in figure legends.

Statistical significance was determined using Student’s t test (between 2 groups) or one-way ANOVA with a tukey post hoc test (multiple comparisons). Individual experiments are noted as different colour shades in the graphs.

## Acknowledgements

The authors thank Tsung-Chang Sung from the Salk Transgenic Core for his assistance generating the DeActs-ER and DeActs-mito constructs. PG was supported by a Queen Elizabeth II Diamond Jubilee Scholarship and a FRQS scholarship. This work was supported by grants from the Natural Sciences and Engineering Research Council of Canada to MG. The Waitt Advanced Biophotonics Core is supported by NIA P30AG068635 (Nathan Shock Center), the Waitt Foundation, and Core Grant application NCI CCSG (CA014195), Grant S10OD030505 (SOM Microscopy Core) and U.M. is supported by NIDCD R01 DC021075-01, NSF NeuroNex Award 2014862, L.I.F.E. Foundation, and the Chan-Zuckerberg Initiative Imaging Scientist Award (https://doi.org/10.37921/694870itnyzk). C.R.S. is supported by NIH 1F32GM137580.

## Data availability

The authors declare that the data supporting the findings of this study are available within the paper and its supplementary information files

---

**Supplemental Figure 1.**
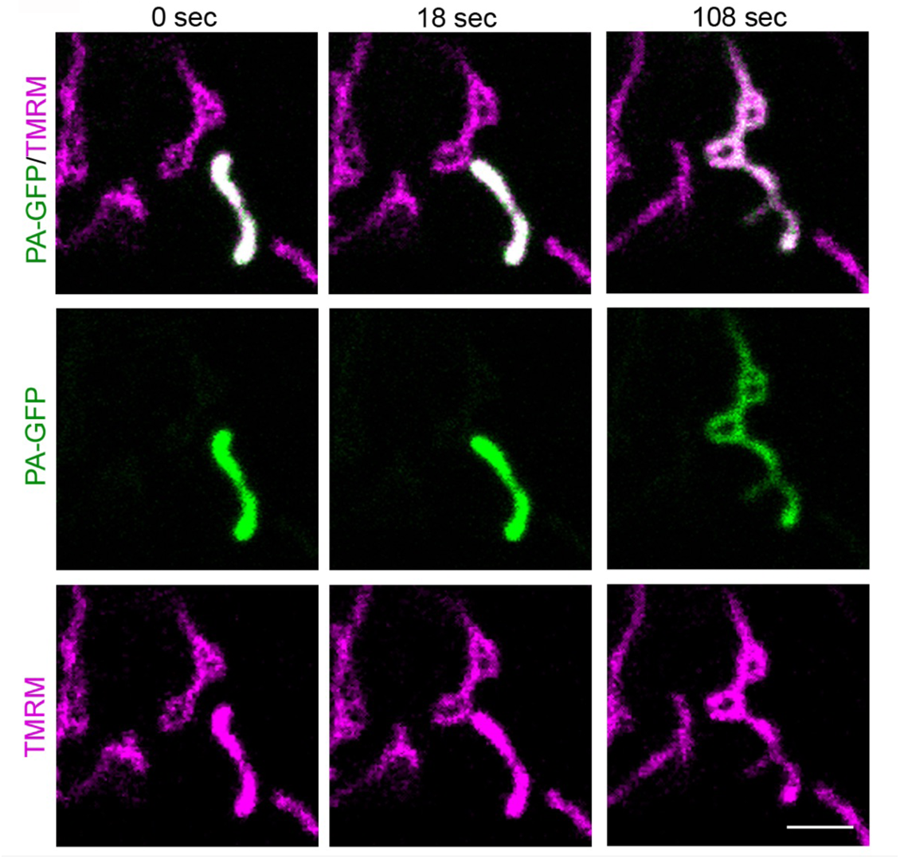
PA-GFP transfer during mitochondrial fusion. (A) Representative images of primary human fibroblasts transfected with PA-GFP (Green) and marked with TMRM (mitochondria, Magenta). The mitochondrion in the lower right corner was activated. Scale bar 2 µm. Related to **Video 1**

**Supplemental Figure 2.**
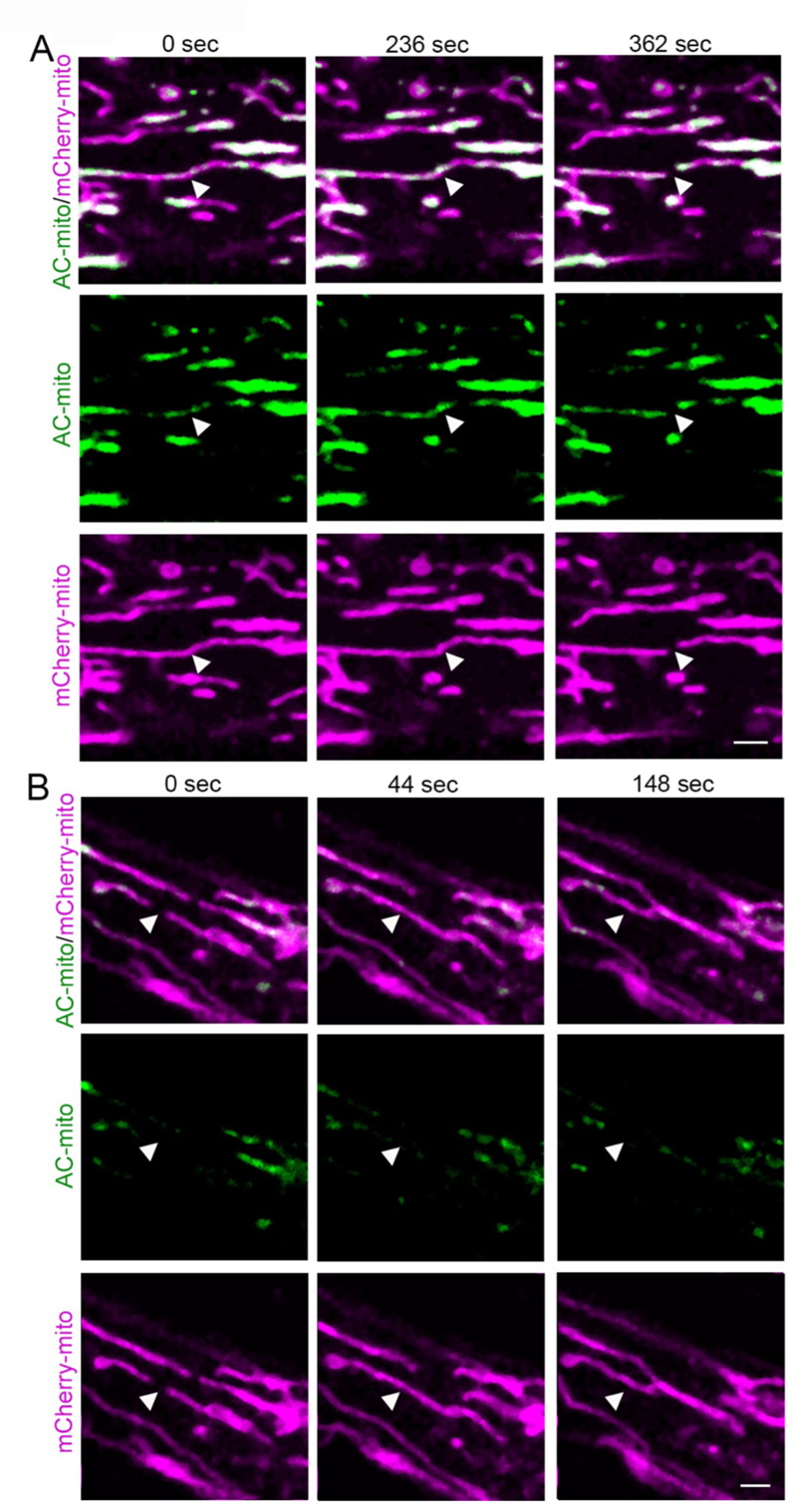
AC-positive fission and fusion events. Representative images of primary fibroblasts transfected with mCherry-mito (mitochondria, magenta) and AC-mito (actin, green). Arrowheads denote a fission event (A) or a tip-to-tip fusion event (B). Scale bar 2 µm. Related to **Video 2** (fission) and **Video 6** (fusion).

**Supplemental Figure 3.**
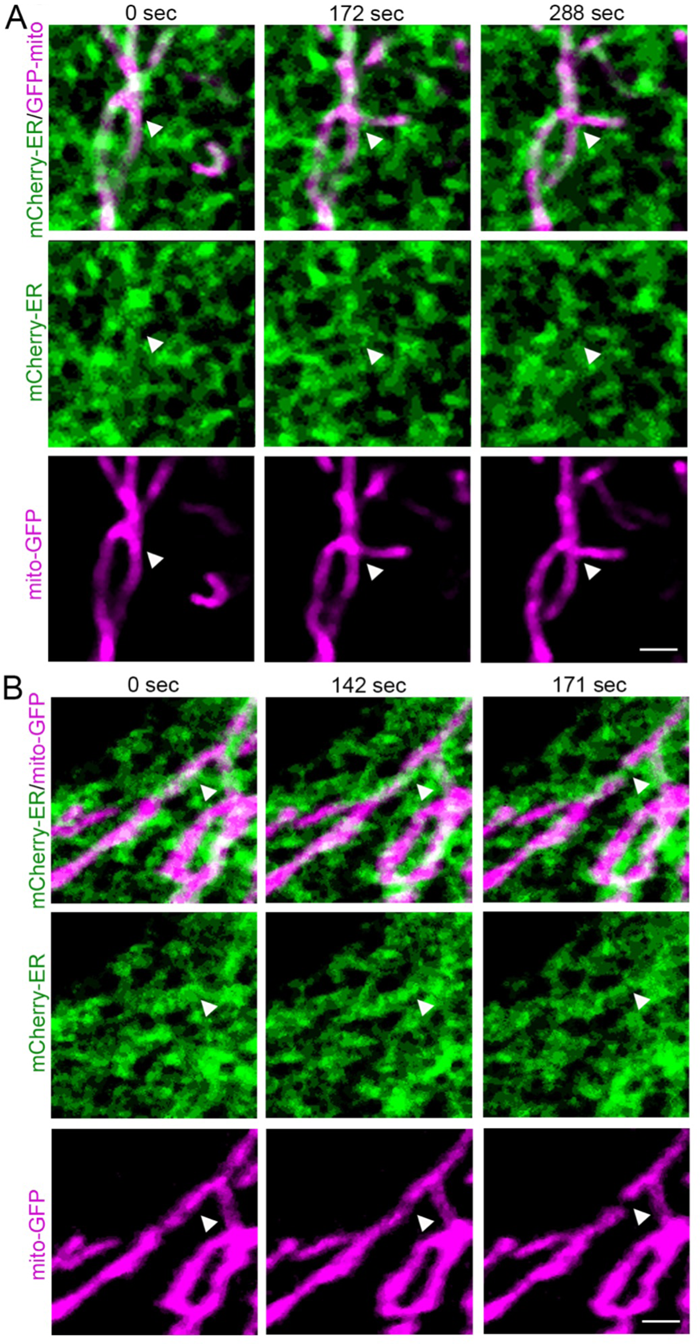
ER-positive fission and fusion events. Representative images of primary fibroblasts transfected with mCherry-ER (ER, Green) and GFP-mito (mitochondria, Magenta). Arrowheads denote a fusion event (A) or a fission event (B). Scale bar 2 µm. Related to **Video 7** (fusion) and **Video 8** (fission).

**Supplemental Figure 4.**
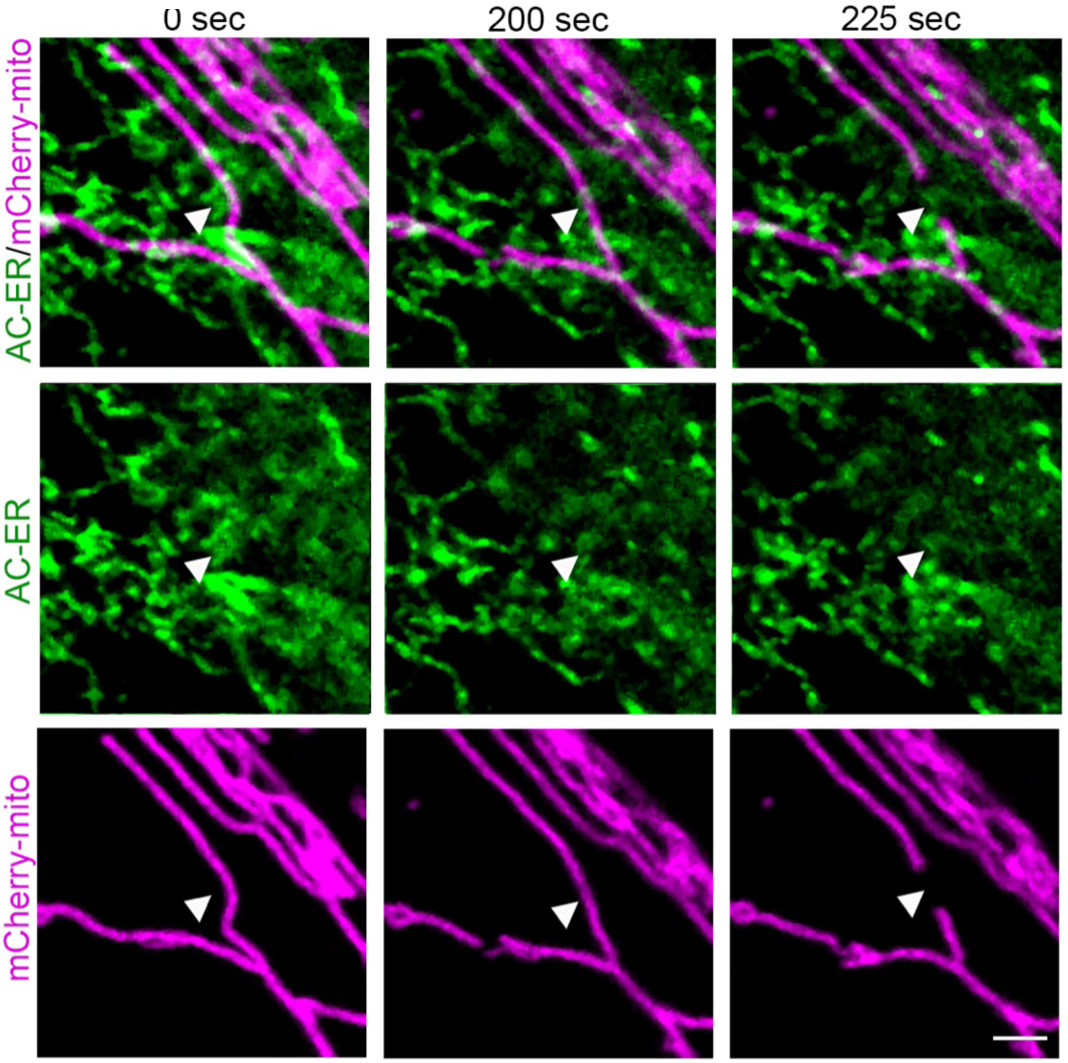
AC-ER-positive fission events. Representative images of primary fibroblasts transfected with AC-ER (Green) and mCherry-mito (mitochondria, Magenta). Arrowheads denote a fission event. Scale bar 2 µm. Related to **Video 10**.

**Supplemental Figure 5.**
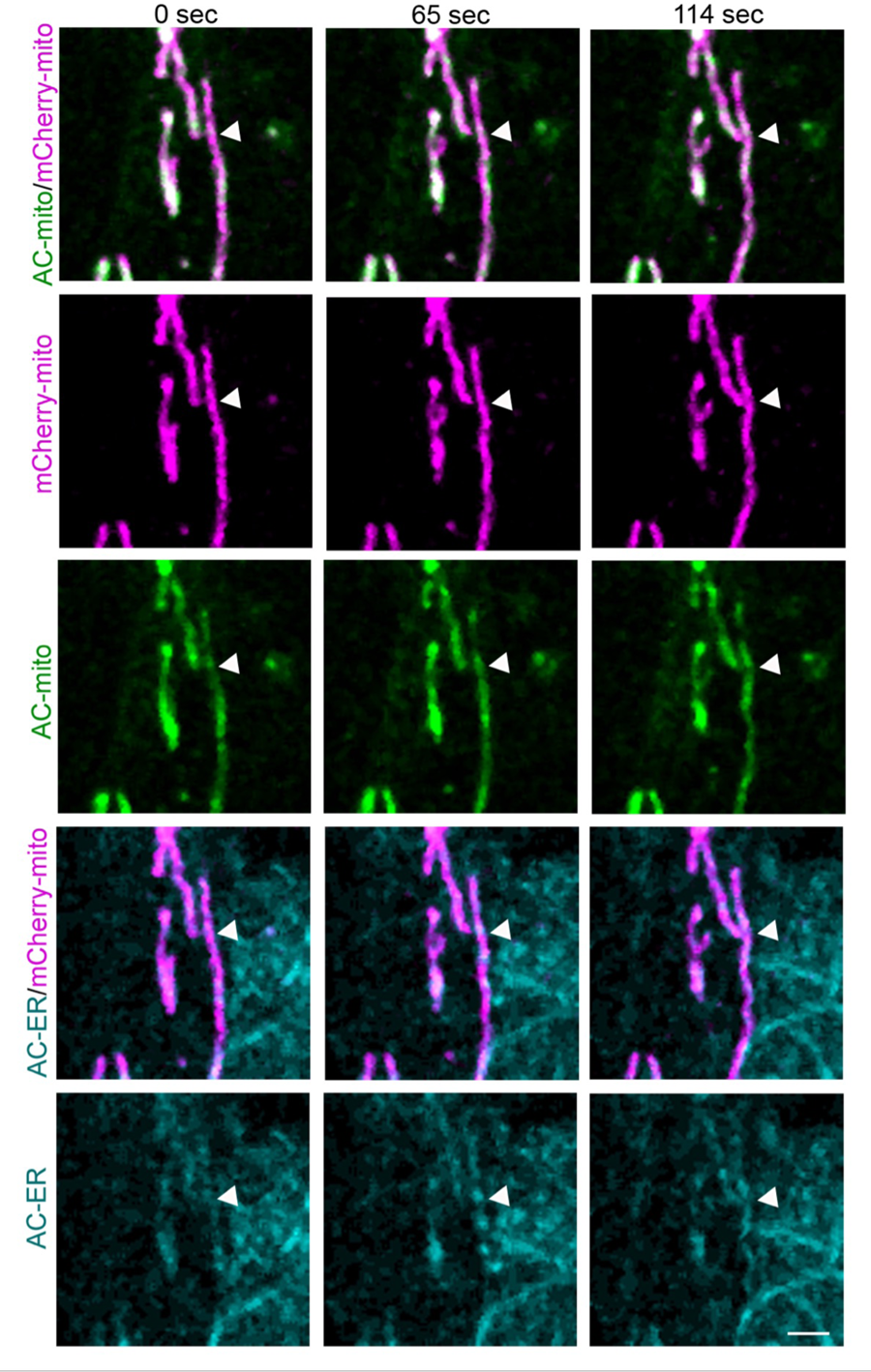
AC-mito accumulates before AC-ER at fusion sites. Representative images of primary fibroblasts transfected with AC-mito (Green), AC-ER (Cian) and mCherry-mito (Magenta). Arrowheads denote a fusion event. Scale bar 2 µm. Related to **Video 11**.

**Supplemental Figure 6.**
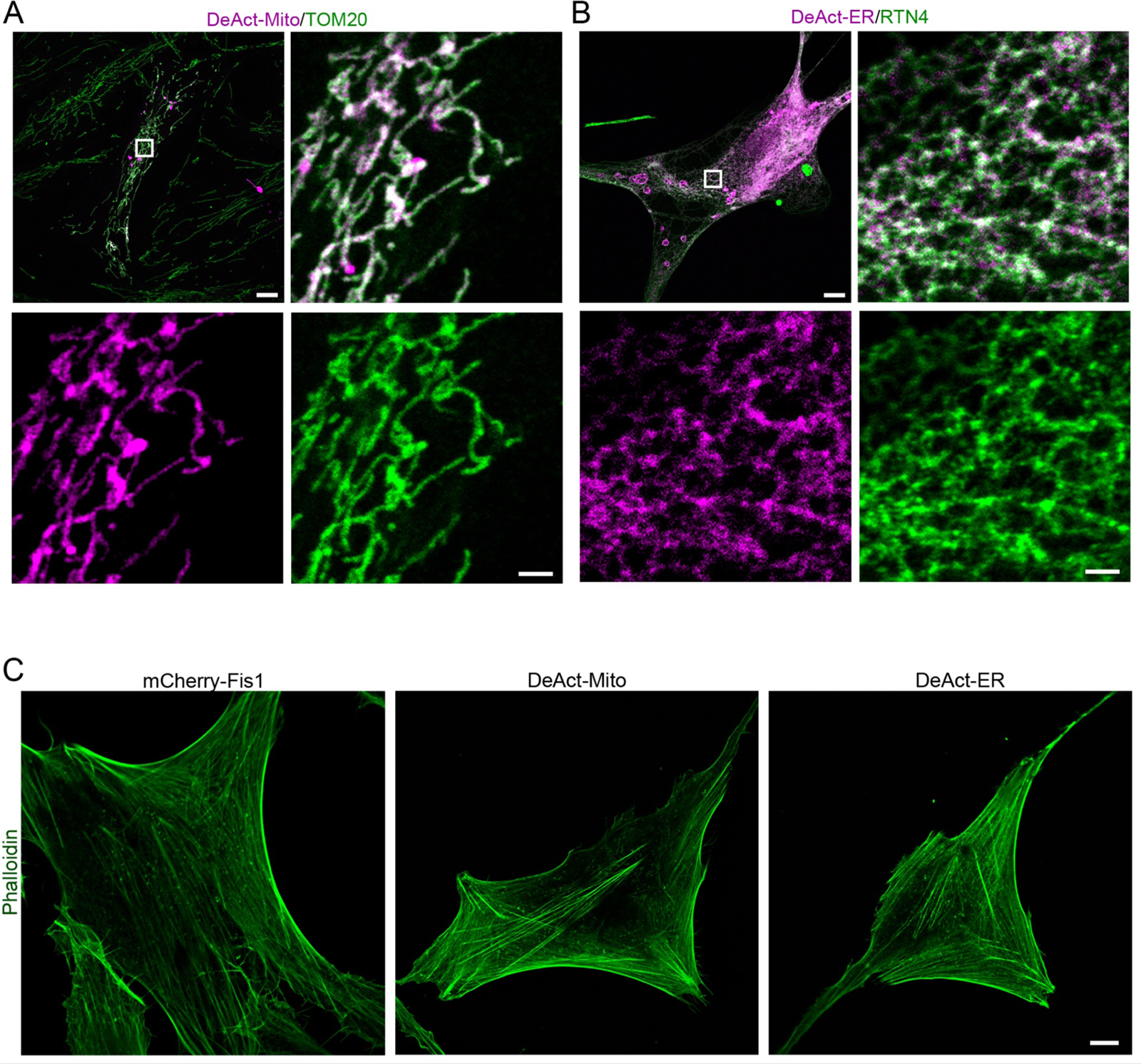
Proper targeting of DeActs. (A) Representative images of primary human fibroblasts transfected with DeAct-mito (magenta) and marked with an antibody against TOM20 (mitochondria, green). The boxed area is shown enlarged below the main image. (B) Representative images of cells transfected with DeAct-ER (magenta) and marked with an antibody against RTN4 (ER, green). The boxed area is shown enlarged below the main image. (C) DeActs do not affect overall actin filaments. Cells were transfected as in (A-B) and stained with phalloidin (actin, green). Scale bar 10 µm, 2 µm for the enlarged images.

**Supplemental Figure 7.**
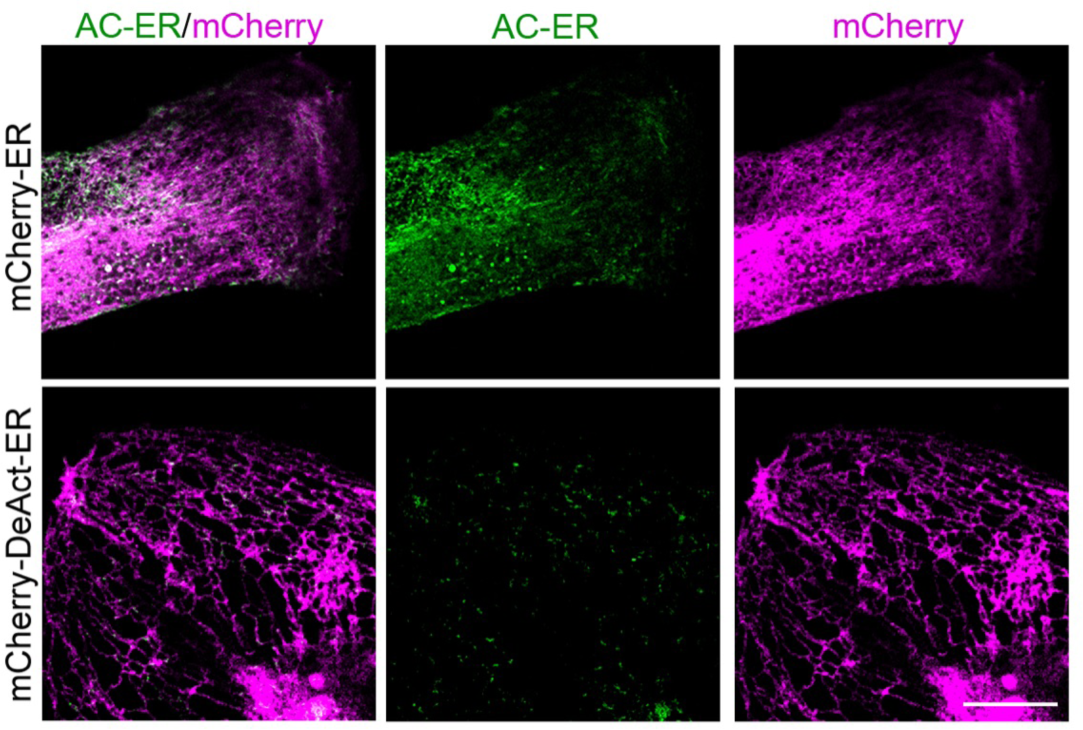
DeAct-ER inhibits the polymerization of ER-associated actin. Representative images showing the loss of AC-ER signal (green) in cells transfected with DeAct--ER (magenta).

**Supplemental Figure 8.**
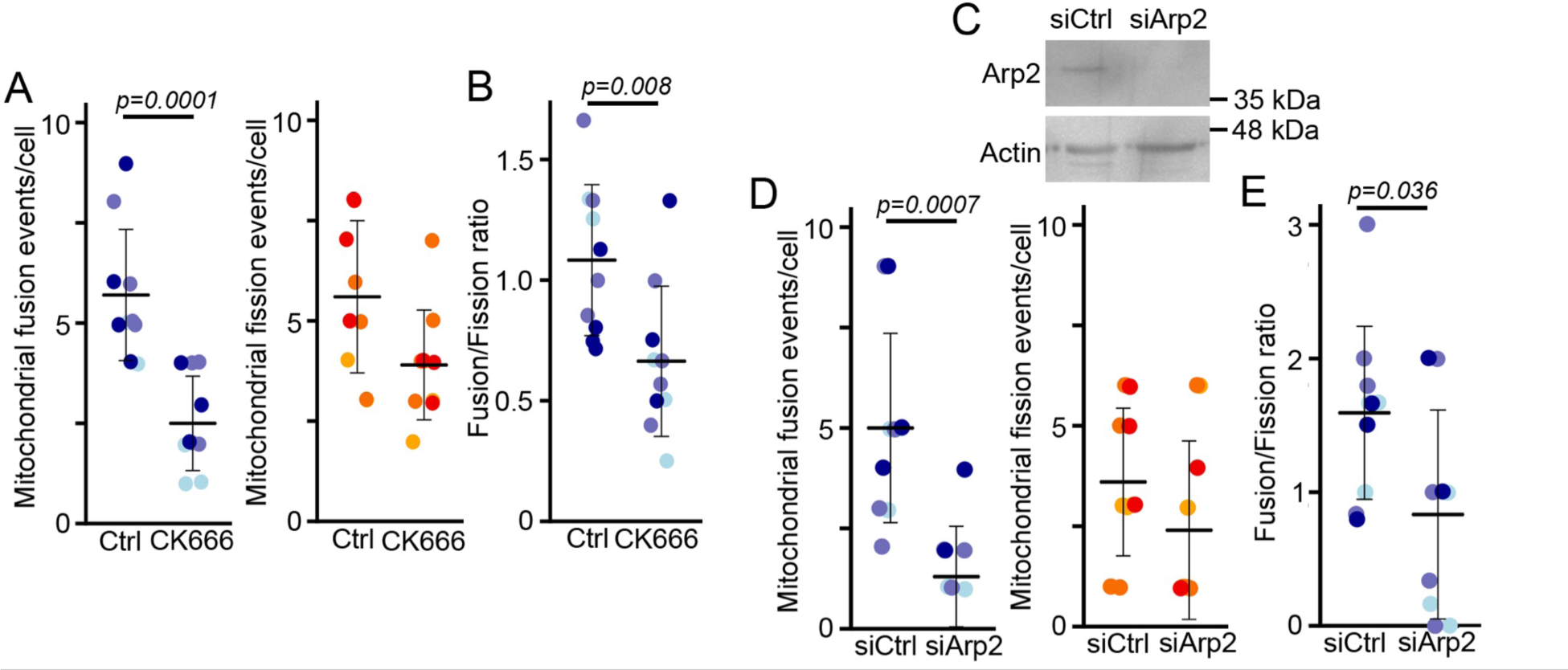
Inhibition of mitochondrial fusion following the inhibition or downregulation of Arp2 in MDA-MB-231 cells. (A) Quantification of the number of fusion (Left, blue) and fission (Middle, orange) events, as well as the fusion/fission ratio (Right) in MDA-MB-231 cells stably expressing mitochondria-targeted GFP and treated as indicated. Each point represents an individual cell, with 10 cells quantified for each condition in 3 independent experiments. Bars show the average ± SD. Two-sided t-test. (B) Western blot showing Arp2 expression in cells treated with a control siRNA (siCtrl) or a siRNA against ARP2 (siARP2). Actin is used as a loading control. (C) Quantification of the number of fusion (Left, blue) and fission (Middle, orange) events, as well as the fusion/fission ratio (Right) in MDA-MB-231 cells stably expressing mitochondria-targeted GFP transfected with siRNAs as in (F). Each point represents an individual cell, with 10 cells quantified for each condition in 3 independent experiments. Bars show the average ± SD. Two-sided t-test.

**Supplemental Figure 9.**
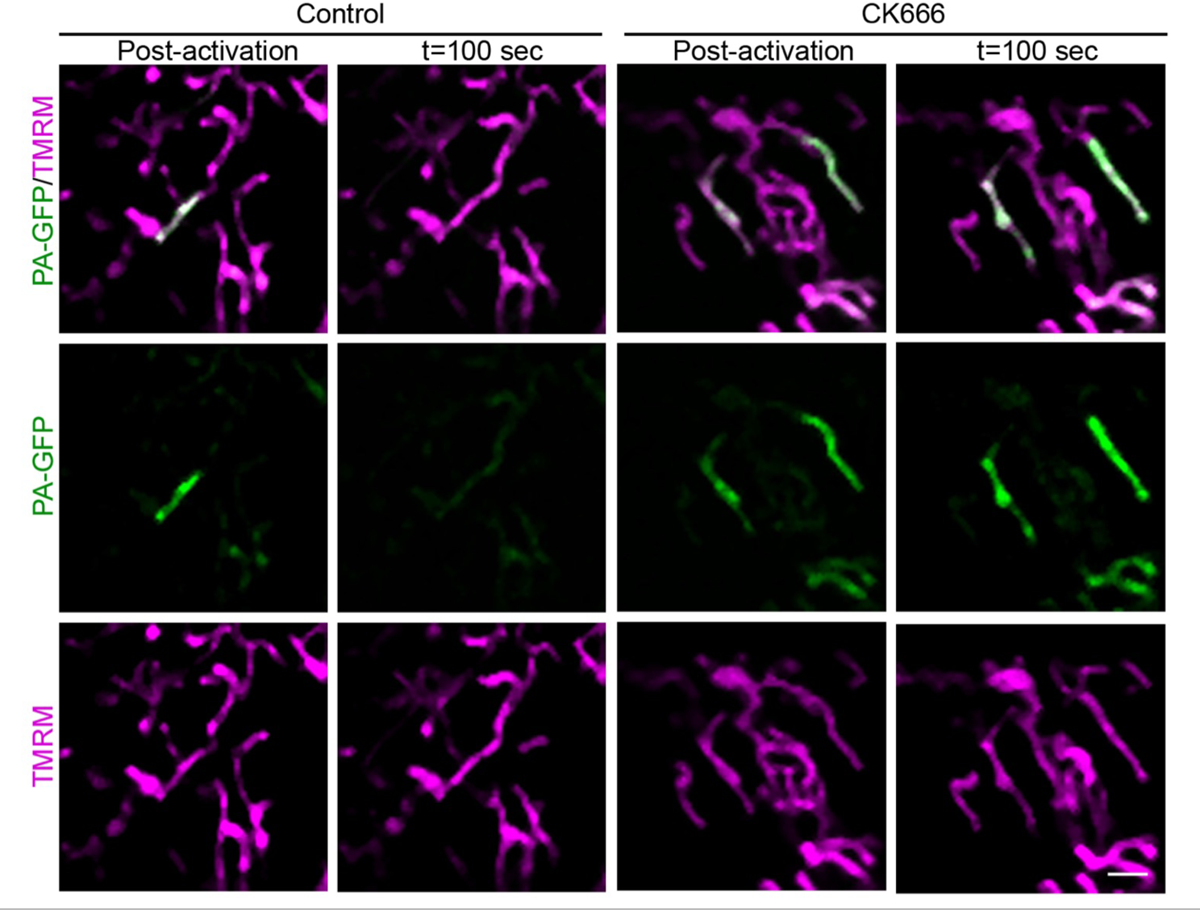
Representative images of the fusion assay. Representative images of human primary fibroblasts transfected with photoactivatable-GFP (PA-GFP) were treated as indicated and imaged immediately after activation (Post-activation) and 100 sec following a fusion event. Scale bar 2 µm.

**Supplemental Figure 10.**
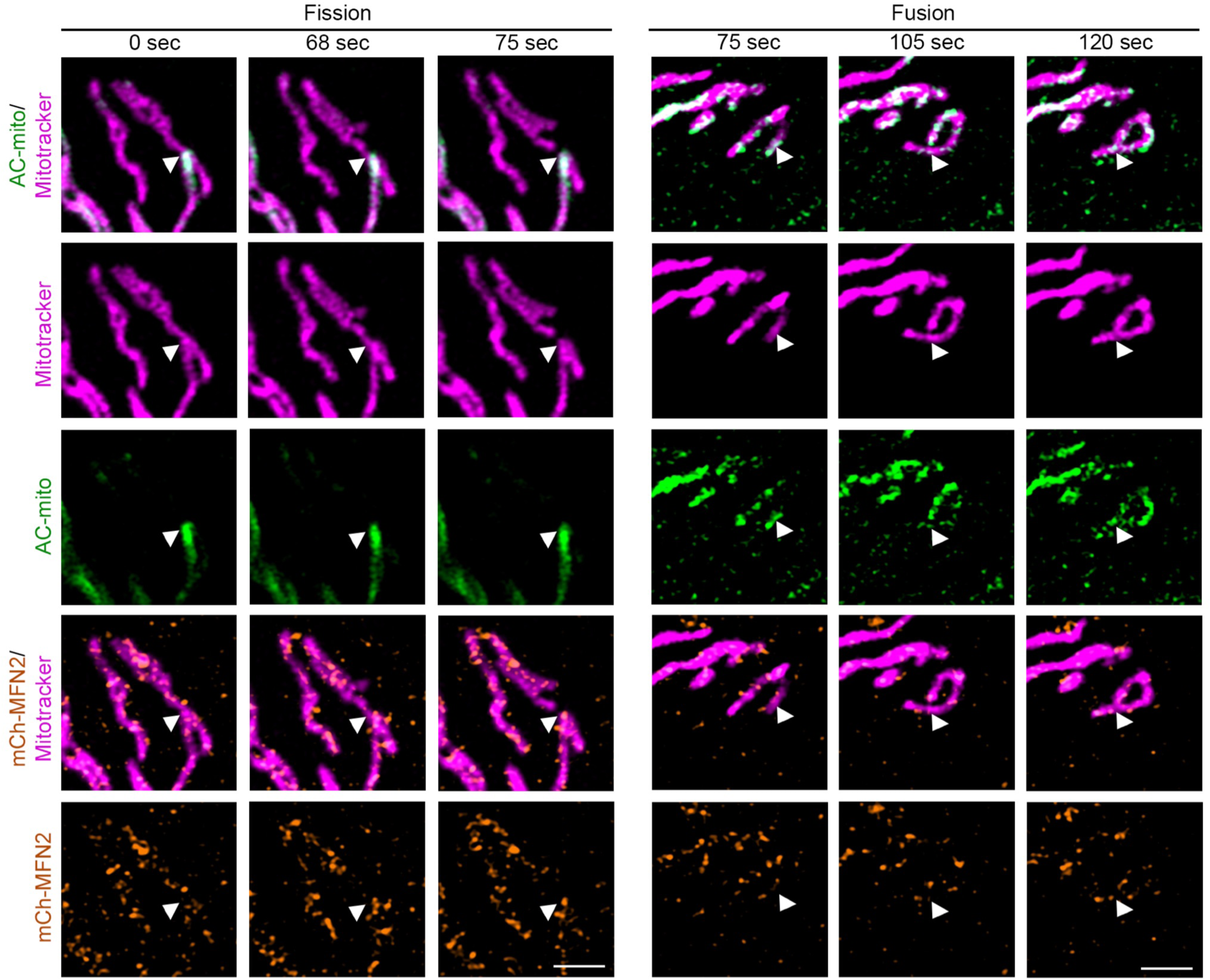
Recruitment of AC-mito and MFN2 at fission and fusion sites. Representative images of primary fibroblasts transfected with mCherry-MFN2 (Orange), AC-mito (Green) and mitochondria marked using mitotracker (Magenta). Arrowheads denote fission (left) and fusion (right) events. Scale bar 2 µm. Related to **Video 12 (fission) and 13 (fusion)**.

**Supplemental Figure 11.**
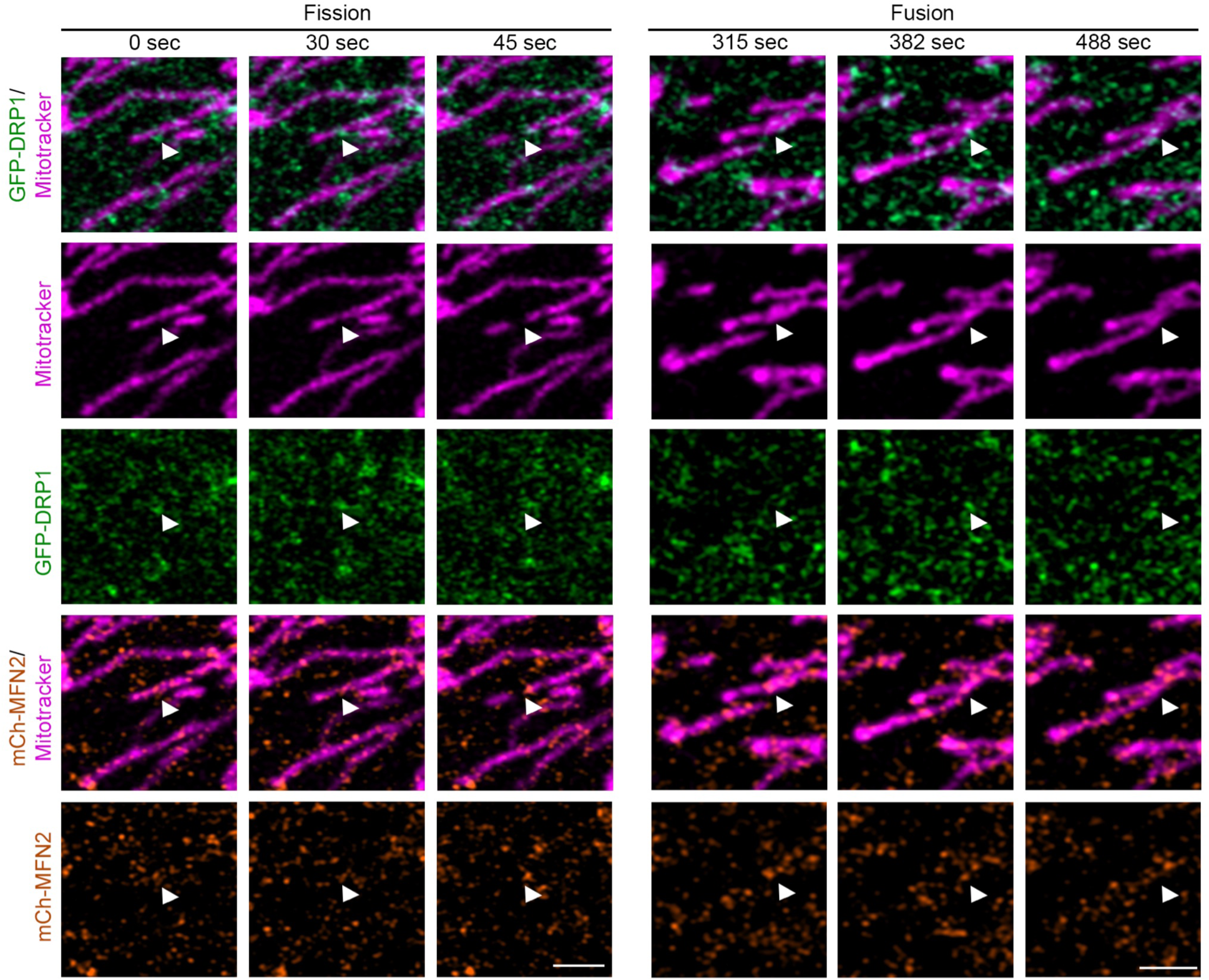
Recruitment of DRP1 and MFN2 at fission and fusion sites. Representative images of primary fibroblasts transfected with mCherry-MFN2 (Orange), GFP-DRP1 (Green) and mitochondria marked using mitotracker (Magenta). Arrowheads denote fission (left) and fusion (right) events. Scale bar 2 µm. Related to **Video 14 (fission) and 15 (fusion)**.

**Supplemental Figure 12.**
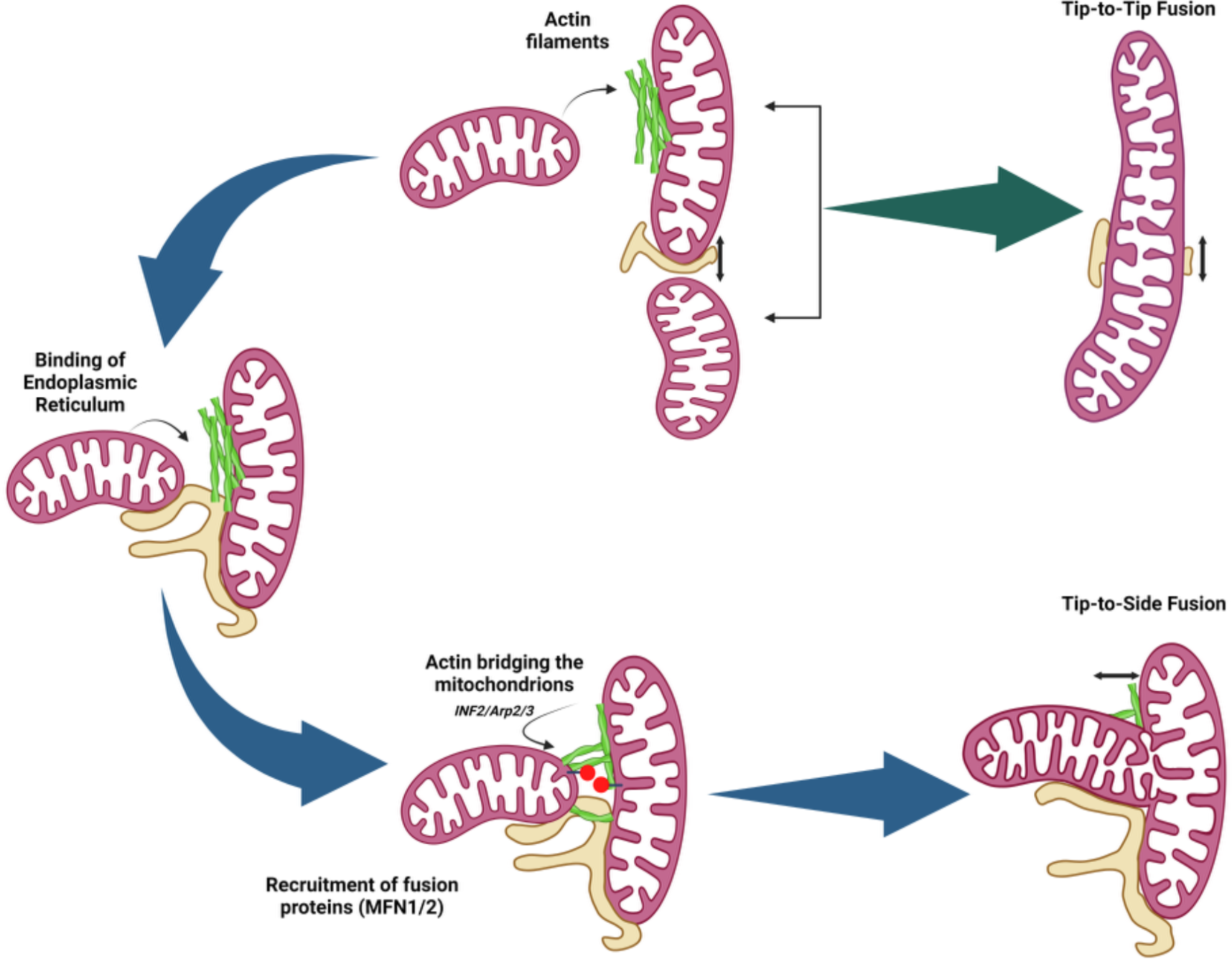
Model for the role of actin in mitochondrial fusion. (A) Tip-to-side fusion requires actin (green). Actin is first polymerised on the stationary mitochondrion by Arp2/3. The ER (yellow-brown) is then recruited to the future fusion site and ER-associated actin is polymerised in an INF2-dependent manner prior to fusion. (B) Tip-to-tip fusion does not require the presence of actin.

## Notes

### Competing Interest Statement

The authors have declared no competing interest.

### Summary of Updates

We have added 1. new lattice-SIM data showing that actin bridges the two fusion mitochondria; 2. new data indicating that actin is recruited to fusion sites prior to MFN2 and DRP1; 3. we have revised and validated all of our assays, including the fusion assay and the validation of DeAct probes.

